# The RNA helicase DHX36/G4R1 modulates *C9orf72* GGGGCC repeat-associated translation

**DOI:** 10.1101/2021.04.25.441260

**Authors:** Yi-Ju Tseng, Siara N. Sandwith, Katelyn M. Green, Antonio E. Chambers, Amy Krans, Heather M. Raimer, Meredith E. Sharlow, Michael A. Reisinger, Adam E. Richardson, Eric D. Routh, Melissa A. Smaldino, Yuh-Hwa Wang, James P. Vaughn, Peter K. Todd, Philip J. Smaldino

**Affiliations:** University of Michigan, Department of Neurology, Ann Arbor, MI; Cellular and Molecular Biology Graduate Program, University of Michigan, Ann Arbor, MI 48109, USA; Ball State University, Department of Biology, Muncie, IN; University of Virginia, Department of Biochemistry and Molecular Genetics, Charlottesville, VA; Lineberger Comprehensive Cancer Center, University of North Carolina at Chapel Hill, Chapel Hill, NC; Nanomedica, Inc., Winston-Salem, NC; Ann Arbor VA Medical Center, Ann Arbor, MI

## Abstract

GGGGCC (G_4_C_2_) hexanucleotide repeat expansions (HRE) in *C9orf72* are the most common genetic cause of amyotrophic lateral sclerosis (ALS) and frontotemporal dementia (FTD). Repeat-associated non-AUG (RAN) translation of this expansion generates toxic proteins that accumulate in patient brains and contribute to disease pathogenesis. The DEAH-Box Helicase 36 (DHX36/G4R1) plays active roles in RNA and DNA G-quadruplex (G4) resolution in cells. As G_4_C_2_ repeats form G4 structures *in vitro*, we sought to determine the impact of manipulating DHX36 expression on repeat transcription and RAN translation. We found that DHX36 depletion suppresses RAN translation from reporter constructs in a repeat length dependent manner while overexpression of DHX36 enhances RAN translation from G_4_C_2_ reporter RNAs. Taken together, these results suggest that DHX36 is active in regulating G_4_C_2_ repeat translation, providing potential implications for therapeutic development in nucleotide repeats expansion disorders.

## INTRODUCTION

A G_4_C_2_ hexanucleotide repeat expansion (HRE) within the first intron of *C9orf72* is a major genetic cause of amyotrophic lateral sclerosis and frontal temporal dementia (C9 FTD/ALS)^1, 2^. Typically, humans have ∼2-28 repeats, while disease associated alleles have >30 and often hundreds to thousands repeats^3, 4^. C9 FTD/ALS represents over 40% of the familial cases and upwards of 10% of the sporadic cases of ALS in European populations^5^. Despite intense research efforts since its discovery in 2011, C9 FTD/ALS remains a progressive and fatal condition without effective treatment^1, 2, 6^.

Both DNA and RNA G_4_C_2_ HRE sequences are prone to folding into G-quadruplex (G4) structures *in vitro*^6–13^. G4 structures are dynamic, nucleic acid secondary structures consisting of an assembly of vertically stacked guanine-tetrad building blocks. G4 structures are stabilized by Hoogsteen hydrogen bonding and monovalent cations, especially K^+^^14–16^. G4 structures have been directly observed in human cells^17–19^, with >700,000 G4 motifs residing throughout the human genome^20, 21^. G4 structures motifs are non-randomly distributed, with enrichment in gene promoters, untranslated regions, and origins of replication, suggesting functional roles in transcription, translation, and replication, respectively^20–24^. Taken together, G4 structures are linked to each of the major toxicities observed in C9 FTD/ALS patient neurons.

The underlying pathogenesis of the G_4_C_2_ HRE involves at least three inter-related pathways, each of which is foundationally linked to aberrant G4 structures. The G_4_C_2_ HRE as DNA impairs mRNA transcription and alters the epigenetics of the *C9orf72* locus, decreasing C9orf72 protein expression^25^. Endogenous C9orf72 protein is important for endosomal trafficking and autophagy in neurons, and its loss is detrimental to neurons and impacts inflammatory pathways relevant to ALS^25^. When transcribed, the resultant G_4_C_2_ mRNA species folds into G4 structures, which coalesce as RNA foci in complex with RNA binding proteins, impairing RNA processing^2, 6^. If transcribed G_4_C_2_ HRE mRNAs reach the cytoplasm, they can serve as a template for repeat-associated non-AUG initiated (RAN) translation. RAN translation from G_4_C_2_ repeat RNA (C9RAN) produces dipeptide repeat proteins (DPRs) that aggregate in proteinaceous inclusions. C9RAN DPRs proteins cause proteotoxic stress and disrupt nucleocytoplasmic transport^13, 26, 27^.

The mechanisms underlying C9RAN remains enigmatic^28^. Initiation can occur at either an upstream near-AUG codon (CUG) or within the repeat itself^29–32^. RNA helicases such as eIF4B, eIF4H and DDX3X play active and selective roles in the translation process, as do the ribosomal accessory protein RPS25^33–36^. RAN translation also demonstrates a selective enhancement in response to cellular stress pathways which activate stress granule formation and suppress global translation through phosphorylation of eIF2α^29–31,37,38^. Consistent with this, modulation of the alternative ternary complex protein eIF2A or PKR expression can alter C9RAN translation.^32, 38^

Given their potentially central role in G_4_C_2_ repeats in C9 FTD/ALS pathogenesis, we explored factors responsible for G4 resolution within cells. One such enzyme, DHX36 (aliases: G4R1 and RHAU), is a member of the DExH-box family of helicases^39^. DHX36 accounts for the majority of the tetramolecular G4 DNA and RNA helicase activity in HeLa cell lysates^40, 41^. DHX36 binds to a diverse array of unimolecular DNA and RNA G4 structures with the tightest affinity of any known G4 structure binding protein and can catalytically unwind these structures in isolation^6, 42–52^. DHX36 associates with thousands of G4-containing DNA and mRNA sequences, facilitating both their transcription and translation^53–55^. Moreover, DHX36 plays an active role in stress granule dynamics, where its loss can trigger spontaneous formation of stress granules and changes in their dissolution after a transient stress exposure^53^. Thus, DHX36 has the potential to influence *C9orf72* transcription and G_4_C_2_-repeat RNA stability, localization, and RAN translation^23, 46^.

Here we find that DHX36 knockdown and knockout selectively suppresses C9RAN translation as well as RAN translation at CGG repeats from reporters in human cells. In contrast, overexpression of WT DHX36, but not a mutant form of DHX36 which lacks helicase activity, enhances RAN translation. These effects are largely translational as we observe suppression of C9RAN translation in an *in vitro* DHX36 KO cell lysate translation system while observing no significant alterations in reporter RNA in response to knockdown or overexpression of DHX36. Loss of DHX36 also precludes stress-dependent upregulation of C9RAN translation consistent with its role in stress granule formation. Taken together, these results suggest modulation of G4 structures at the RNA level by candidate G4 helicases such as DHX36 impact G_4_C_2_ repeat expansion translation implicated in C9 FTD/ALS.

## RESULTS

### DHX36 directly binds C9-repeat G4 DNA *in vitro*

To determine if DHX36 directly binds to C9 G4 DNA structures, we performed electrophoretic mobility shift assays (EMSAs) with a DNA oligonucleotide composed of five G_4_C_2_-repeats with a 3’ unstructured tail shown to be require for G4 binding^49, 56^ (referred to hereafter as “(G_4_C_2_)_5_-DNA”). (G_4_C_2_)_5_-DNA was folded into G4 structures by heating and cooling in the presence of KCl. As a negative control of non-G4 DNA, (G_4_C_2_)_5_-DNA was heated and cooled in the absence of KCl. G4 (G_4_C_2_)_5_-DNA and non- G4 (G_4_C_2_)_5_-DNA was incubated with purified recombinant DHX36 (rDHX36) under non-catalytic conditions (-ATP, +EDTA) so that binding could be visualized on a gel **(Figure 1A, C)**. As an additional control, this was repeated with scrambled C9-repeat DNA, where the C9-repeat sequence was rearranged as to prevent G4 structure formation. **(Figure 1B-C)**. Following incubation, the samples were subjected to non-denaturing polyacrylamide gel electrophoresis (PAGE). In the absence of KCl, a single band is observed for (G_4_C_2_)_5_-DNA. When KCl is added, slower migrating bands are observed, consistent with the formation of G4 structures. Incubation of (G_4_C_2_)_5_-DNA with rDHX36 resulted in a shift of the DNA to the upper region of the gel indicating direct binding. Notably, some unstructured (G_4_C_2_)_5_-DNA is present in the KCl-containing reactions and is not bound by DHX36, further suggesting selective binding to G4 structure. In the absence of KCl (i.e. non-G4 conditions), binding of rDHX36 to (G_4_C_2_)_5_-DNA is substantially reduced. Furthermore, scrambled (G_4_C_2_)_5_-DNA does not form KCl-dependent higher-ordered structures and is not a strong binding substrate for rDHX36 even in the presence of KCl. Taken together, these data suggest that DHX36 directly binds to C9 HRE DNA in a G4-dependent manner *in vitro*.

**Figure 1:**
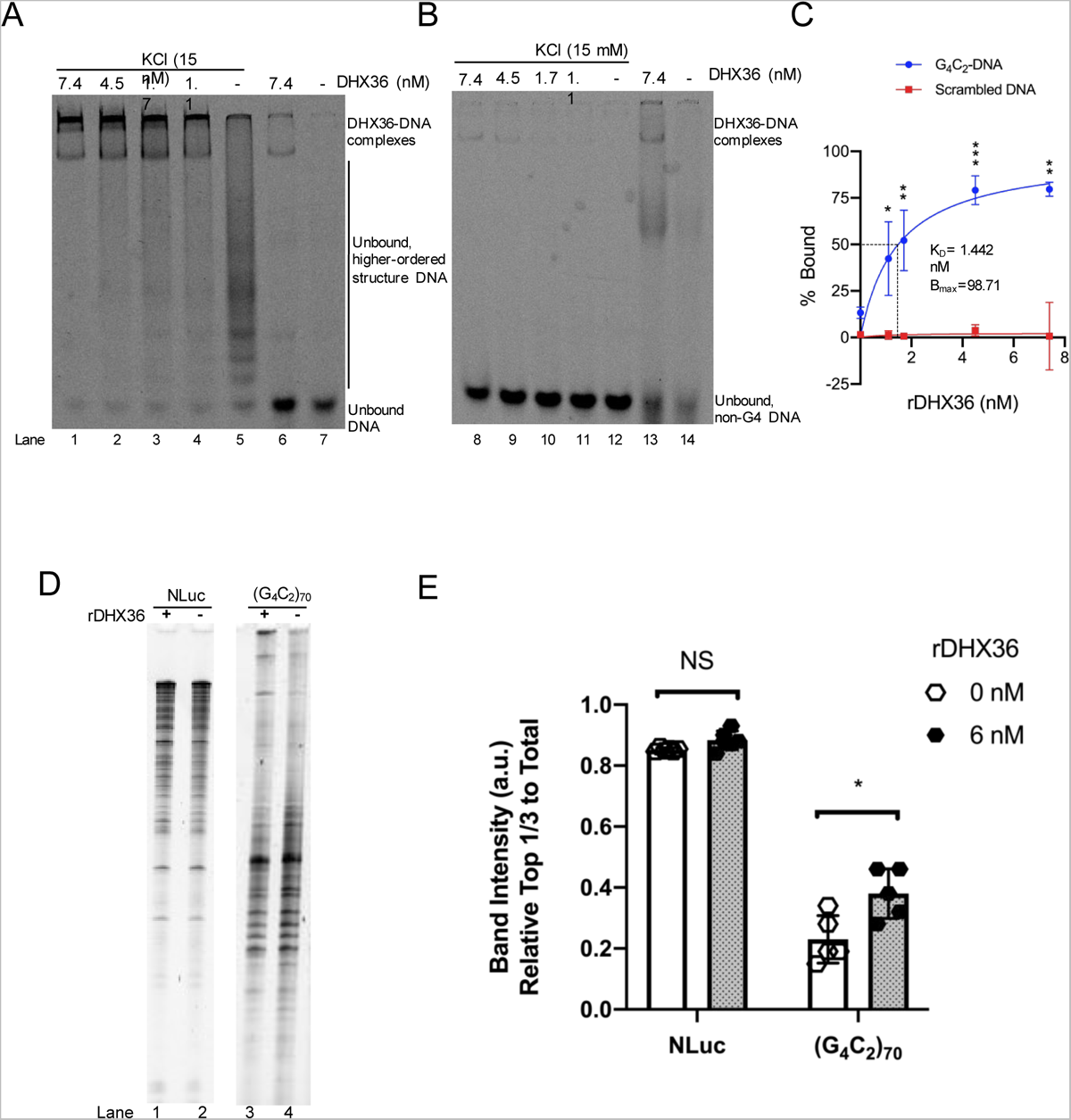
DHX36 binds and enhances transcription of C9-DNA *in vitro.* (A) Representative electrophoretic mobility shift assay (EMSA) image. *C9* repeat DNA oligonucleotides were heated and cooled in the presence (lanes 1-5) and absence (lanes 6-7) of KCl to induce or prevent G4 formation, respectively. DNA was incubated with increasing concentrations of recombinant DHX36, analyzed with non-denaturing PAGE, and imaged. (B) Representative EMSA image. Scrambled control DNA oligonucleotides were heated and cooled in the presence (lanes 1-5) and absence (lanes 6-7) of KCl. DNA was incubated with increasing concentrations of recombinant DHX36, analyzed with non-denaturing PAGE, and imaged. (C) Densiometric quantification of panels A and B. The percent bound for each lane was graphed versus the concentration of DHX36. Data in are presented as mean ±SD, n=3. Multiple t-tests for each concentration of protein, *p=<0.05, **p<0.01, ***p<0.001. (D) T7 polymerase transcript products generated from equal amounts of linearized nanoLuciferase (lanes 1-2) or (G_4_C_2_)_70_ plasmids (lanes 3-4) resolved by denaturing PAGE. The gel was stained with SYBR gold nucleic acid stain and imaged. (E) Densiometric quantification of panel D. All signals were first subtracted by the background. Then signal from the top third of the gel was divided by the total signal per lane. Data are presented as ±SD, n=5(NLuc)-6(G_4_C_2_)_70_. Two-tailed pair t-test, N.S. = not-significant, \**P* < 0.05.

### DHX36 enhances transcription of C9-repeat DNA *in vitro*

G4 DNA structures impede the transcription of C9-repeat RNA and T7 elongation ^6^. Given that DHX36 is a helicase that resolves G4 structures, we hypothesized that DHX36 might facilitate the transcription of C9-repeat RNA. To test this, we performed an *in vitro* transcription assay with a plasmid containing 70 G_4_C_2_ repeats (pCR8- (G_4_C_2_)_70_) driven by a T7 RNA polymerase reporter ^27^. We incubated the plasmid with T7 polymerase in the presence and absence of rDHX36. A T7 plasmid containing a nano luciferase (NLuc) gene was used as a non-G4 control. The resulting RNA transcripts were subjected to denaturing urea gel electrophoresis. We found that rDHX36 significantly increased the length of RNA transcripts yielded from G_4_C_2_ repeat DNA, but not from NLuc **(Figure 1D-E)**. However, the total RNA generated from G_4_C_2_ repeat DNA and NLuc DNA was not significantly different between rDHX36 and control reactions **(Supplemental Figure 1)**. These data suggest that rDHX36 facilitates efficient and complete *in vitro* transcription of G4 C9-repeat sequences by T7 RNA polymerase, but may not impact its overall production.

### DHX36 depletion modifies C9RAN translation

We next evaluated the impact of altering DHX36 expression on C9RAN translation. To accomplish this, we utilized previously described C9RAN translation-specific nanoluciferase reporters (C9-NLuc)^29^. These reporters include 70 G_4_C_2_ repeats in the context of the first *C9orf72* intron. This sequence is inserted 5’ to a nanoluciferase (NLuc) whose start codon is mutated to GGG and with a 3x FLAG tag fused to its carboxyl terminus **(Figure 2A)**. Single base pair insertions between the repeat and NLuc allow for evaluation of translation in all three reading frames. An AUG-initiated NLuc serves as a positive control for canonical translation. An AUG initiated firefly luciferase (AUG-FF) is included as a transfection control_29_.

**Figure 2:**
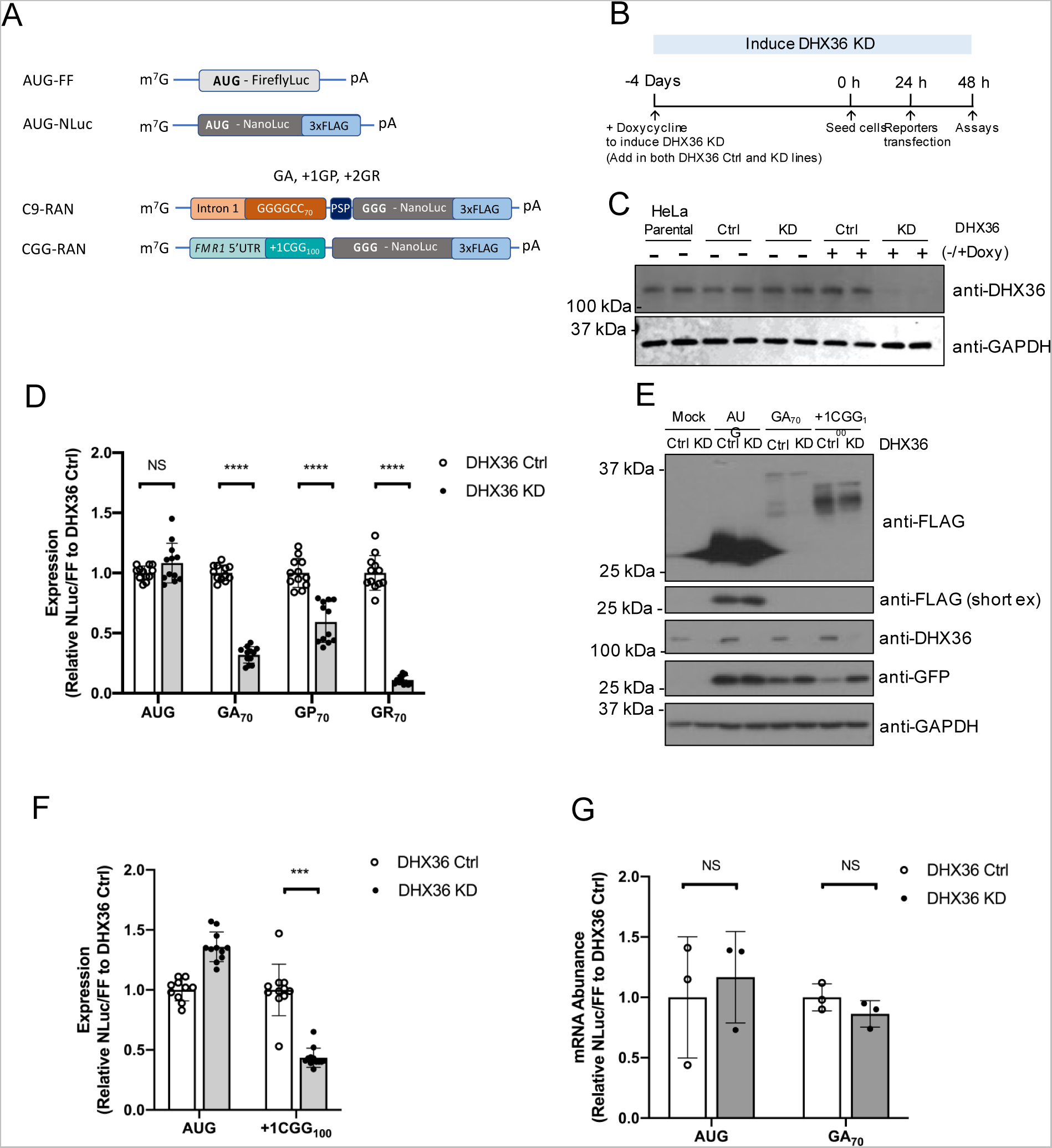
The effect of DHX36 knockdown on C9-RNA and C9RAN reporter expression. (A) Schematic of AUG-FF, AUG-NLuc control, C9-RAN and CGG-RAN luciferase reporters. (B) Experimental timeline for doxycycline treatment and reporter transfection. (C) Immunoblots detecting DHX36 in parental, Ctrl and DHX36 KD HeLa cells with and without doxycycline treatment. (D) Relative expression of AUG and C9- RAN translation in GA (+0), GP (+1), and GR (+2) frames with 70 repeats between Ctrl and DHX36 KD HeLa cells. NLuc signal were normalized to AUG-FFluc translation. (E) Immunoblot of RAN translation products from 70 repeats of G_4_C_2_ in GA frame and 100 repeats of CGG in +1 reading frame in Ctrl and DHX36 KD HeLa cells. GFP was blotted as transfection control and GAPDH was blotted as loading control. (F) Expression of +1CGG_100_ RAN translation reporters measured by luciferase assay. NLuc signals were normalized to AUG-FFluc signals to compare between Ctrl and DHX36 KD HeLa cells. (G) Abundance of NLuc mRNA from AUG and GA_70_ in DHX36 Ctrl and KD HeLa cells. NLuc mRNA were normalized to FF mRNA and compared to DHX36 Ctrl. Data in (D) and (F) are represented as mean ±SD, n=9-12. Data in (G) are mean ±SD, n=3. Two- tailed Student’s t-test with Bonferroni and Welch’s correction, N.S. = not-significant, \**P* < 0.05; \*\**P* < 0.01; \*\*\**P* < 0.001; \*\*\*\**P* < 0.0001.

To study the effects of loss of DHX36, we used a previously described stable and inducible knockdown (DHX36 KD) HeLa cell line_41,57_ **(Figure 2B, Supplemental Figure 2A-E)**. Treatment of these cells with doxycycline for 96 hrs significantly reduced DHX36 expression as measured by immunoblot **(Figure 2C)**. Comparing between control and DHX36 KD cells with transiently transfected C9-NLuc reporters, DHX36 KD selectively decreased C9RAN translation in the GA (+0), GP (+1) and GR (+2) reading frames, relative to AUG-NLuc when normalized to AUG-FF as transfection control **(Figure 2D, Supplemental Figure 2F)**. C9RAN translation in the GA and GR reading frames was also selectively decreased in the DHX36 KD line with doxycycline induction when compared to the DMSO vehicle treated cells, suggesting that the effect was DHX36- specific **(Supplemental Figure 3A).** Either no significant transcript bias or an opposite production bias favoring DHX36 KD was observed for AUG transcripts **(Figure 2D and F)**. To confirm these findings using an orthogonal readout, we performed immunoblots to detect FLAG signal on lysates from both control and DHX36 KD cells. As we had observed in our luciferase assays, DHX36 KD led to a significant decrease in the GA C9RAN-NLuc protein without impairing AUG translation of NLuc from a separate reporter (**Figure 2E**).

To determine if loss of DHX36 might have broader effects on protein translation in these cells, we performed a SUnSET assay, which measures puromycin incorporation into nascent proteins_58_. Treatment with puromycin for 10 minutes led to a smear of proteins detectible by puromycin immunoblot. There was no difference between DHX36 Ctrl and KD cells in this assay (**Supplemental Figure 2D-E**), suggesting that rates of global translation is not demonstrably affected by knockdown of DHX36 in these cell lines.

### DHX36 depletion impairs RAN translation from CGG repeats

RAN translation occurs at multiple different GC rich repeat sequences, some of which are capable of forming G4 structures and some of which are less likely to form such structures. We therefore evaluated whether DHX36 KD impacts RAN translation at these other repeats. Expansion of a transcribed CGG repeats in the 5’UTR of *FMR1* causes Fragile X-associated Tremor/Ataxia Syndrome (FXTAS)_59,60_. RAN translation from this repeat in the +1 reading frame generates a polyglycine protein (FMRpolyG) that accumulates within inclusions in patient brains and model systems_59,61–64_. This repeat is capable of forming either a hairpin structure or a G4 structure *in vitro*_65–68_. Using a NLuc reporter with 100 repeats (+1CGG_100_)^69^, we observed that KD of DHX36 significantly suppressed CGG RAN translation of FMRpolyG on a scale comparable to that of C9RAN reporters **(Figure 2F)**. This decrease in CGG RAN translation was also evident by immunoblot **(Figure 2E)**.

We next measured reporter mRNA levels in transfected Ctrl and DHX36 KD cells. Surprisingly, we observed only a small decrease in mRNA production that was not statistically significant **(Figure 2G)**. In parallel, we also evaluated the impact of DHX36 overexpression on reporter expression. As with KD, overexpression of DHX36 did not significantly impact steady state reporter RNA expression in HeLa Cells **(Supplemental Figure 5)**. These data suggest that the suppression of RAN translation in DHX36 KD cells is most likely a post-transcriptional event.

### DHX36 KO impairs in-cell and *in vitro* C9RAN translation

As a second assay system in which to study the effect of DHX36, we generated a DHX36 stable knockout (DHX36 KO) Jurkat cell line using a CRISPR/Cas9 targeting approach. Wildtype Jurkat cells (WT) had no mutations at the DHX36 locus (INDEL: 0/0) while DHX36 KO Jurkat cells (DHX36 KO) had single allele KO disruption on one allele and a 6bp insertion on the other allele at the target site (INDEL: +5/+6) **(Supplemental Figure 4A-B)**. Western blot analysis showed elimination of full length DHX36 protein in Jurkat DHX36 KO cell lines **(Figure 3A)**. Jurkat DHX36 KO cells exhibited impaired RAN translation across all three potential G_4_C_2_ reading frames, similar to what we observed in HeLa DHX36 KD cells **(Figure 3B)**.

**Figure 3:**
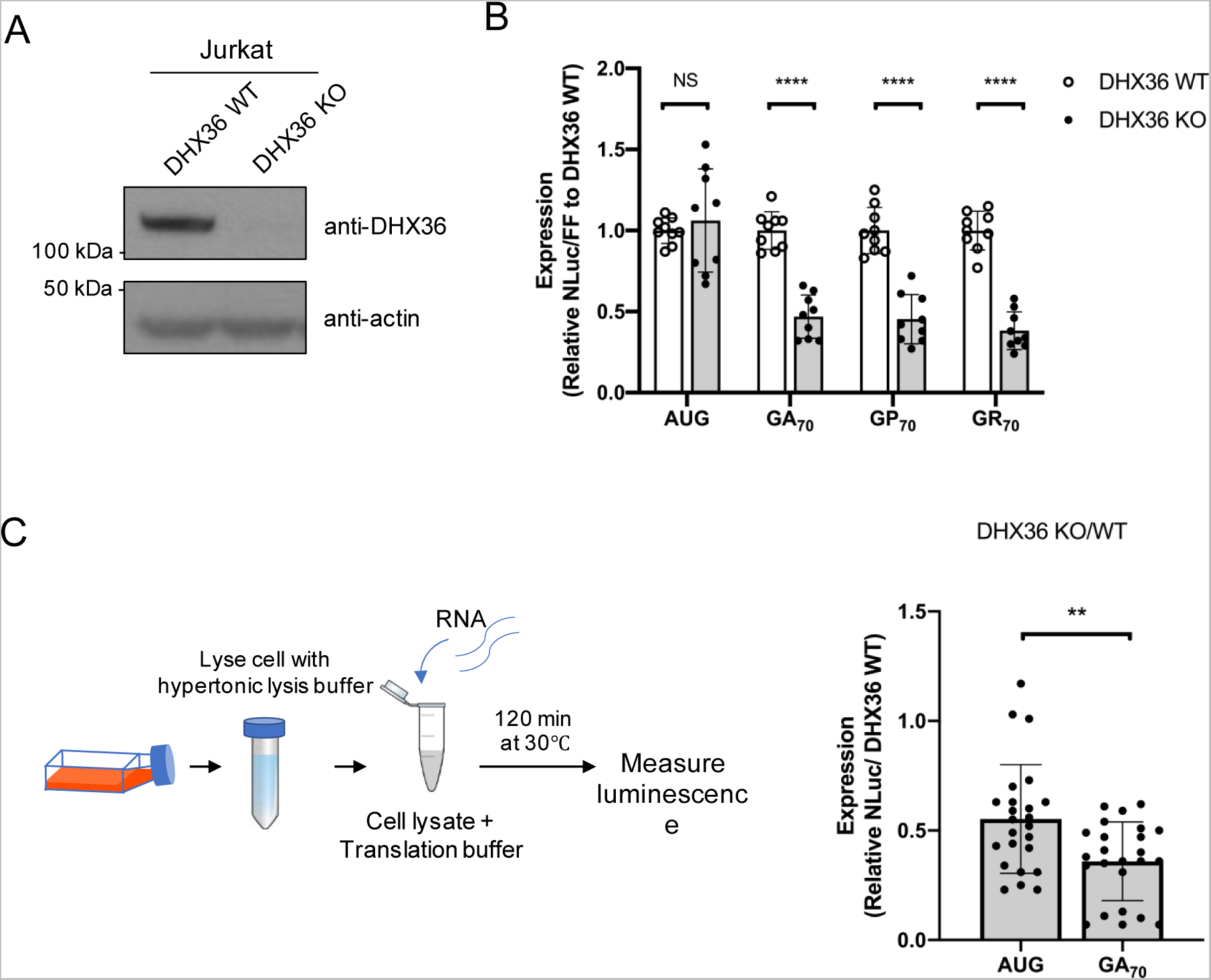
C9RAN reporter expression in DHX36 knockout Jurkat cell lines and *in vitro* cell lysates. (A) Immunoblots to DHX36 from WT and DHX36 KO Jurkat cells. (B) Relative expression of AUG and C9-RAN translation from GA(+0), GP(+1), and GR(+2) reading frames in WT and DHX36 KO Jurkat cells. NLuc signal were normalized to AUG-FF and compared between WT and DHX36 KO Jurkat cells. Data are represented as mean ±SD, n=9. (C) *In vitro* translation using lysates derived from DHX36 WT and KO Jurkat cells for AUG-NLuc RNA and C9-RAN in GA frame RNA. NLuc signals were normalized to signal from DHX36 WT. Data are represented as mean ±SD, n=24. Two- tailed Student’s t-test with Bonferroni and Welch’s correction, \**P* < 0.05; \*\*\**P* < 0.001; \*\*\*\**P* < 0.0001.

The effects of DHX36KD on RAN translation product generation could theoretically be elicited by changes in RNA or protein stability or by actively impacting protein translation. To investigate this question, we utilized an *in vitro* translation assay using lysates derived from DHX36 WT or KO Jurkat cells. Previous studies in similar conditions demonstrated that we could accurately measure C9RAN translation in this context and that production from our RAN reporters is not dependent on mRNA or reporter stability^29, 69^. We harvested Jurkat DHX36 WT and KO cell lysates, added AUG or G_4_C_2_ repeat RNA in GA (+0) frame and *in vitro* translated for 2 hrs **(Figure 4C, Supplemental Figure 4C)**. AUG-NLuc translation from DHX36 KO lysates was consistently lower than that from WT Jurkat lysates. However, this effect was much larger for GA-NLuc reporters, which exhibited 36% as efficient a translation in DHX36 KO lysates compared to WT lysates over >4 independent experiments **(Supplemental Figure 4C)**. Together, these results suggest that loss of DHX36 suppresses RAN translation from G_4_C_2_-repeats in multiple reading frames of G_4_C_2_-repeats and is mainly acting at the level of translation.

**Figure 4:**
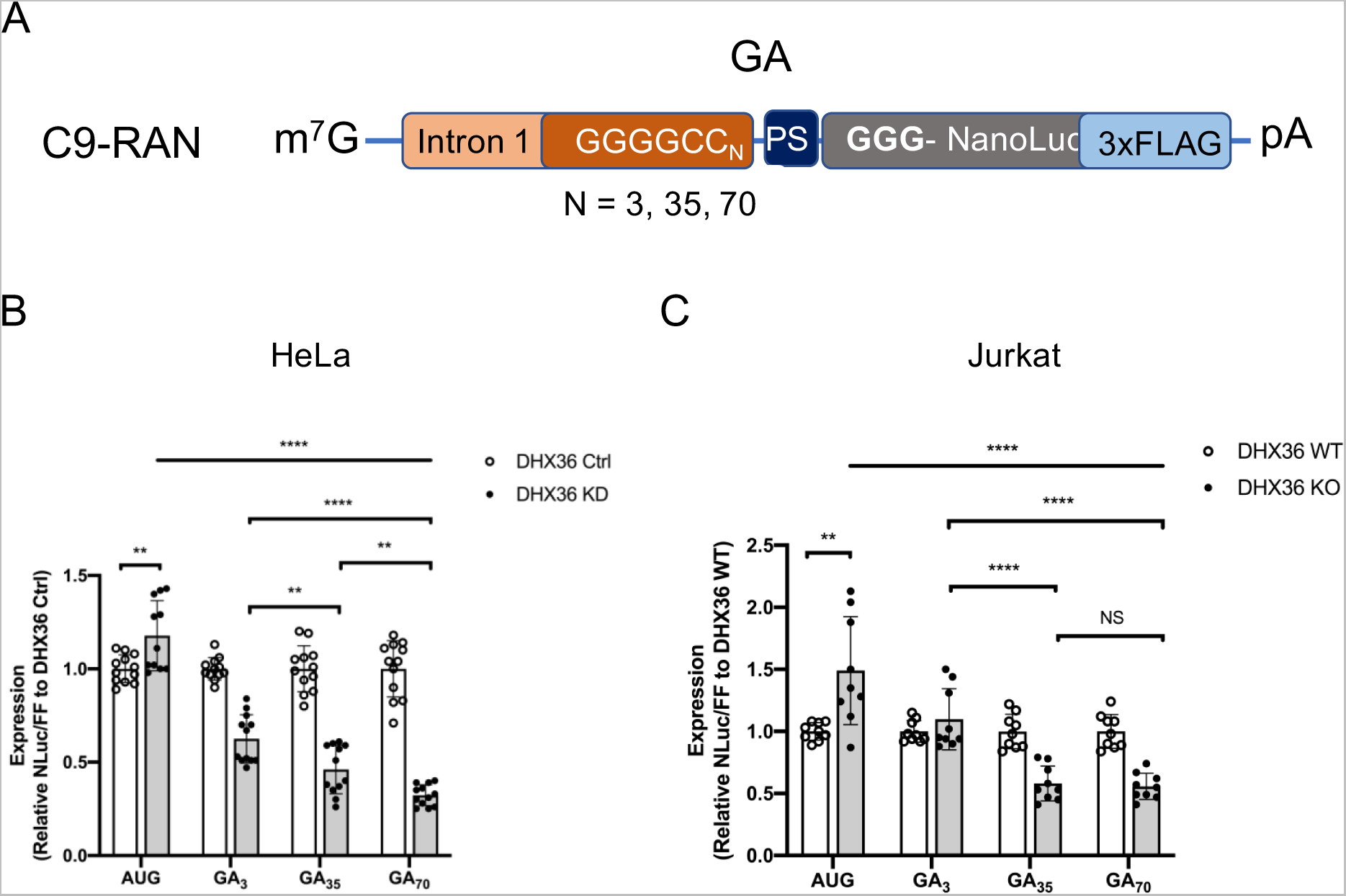
The effect of decreased DHX36 on C9RAN reporter expression is G_4_C_2_ repeat length dependent. (A) Schematic of previously published luciferase reporters of C9-RAN in GA frame harboring different repeat sizes. (B)-(C) Relative expression of AUG and C9-RAN translation from GA frames with 3, 35, and 70 repeats in Ctrl and DHX36 KD HeLa cells (B), and DHX36 WT and KO Jurkat cells. NLuc signal were normalized to AUG-FF. Data are represented as mean ±SD, n=9-12. One-way ANOVA were performed to compare the statistical differences between repeat length in DHX36 KD or KO cell lines. Two-tailed Student’s t-test with Bonferroni and Welch’s correction were then perform to confirm the differences between multiple comparison, \**P* < 0.05; \*\**P* < 0.01; \*\*\*\**P* < 0.0001.

### The effect of DHX36 on C9RAN translation is dependent on repeat length

Translation of C9RAN reporters in the GA reading frame initiates primarily from an upstream CUG start codon that supports translation even at small repeat sizes^29, 31, 32, 70^. If DHX36 contributes to RAN translation by resolving G4 structures, then we would predict that the loss of DHX36 would selectively reduce translation for transcripts with larger repeats. In HeLa cells, expression of C9 GA frame reporters with 3x, 35x, and 70x G_4_C_2_ repeats was selectively suppressed by DHX36 loss at the larger repeat sizes **(Figure 4A-B, Supplemental Figure 3B)**. Similar results were observed in Jurkat DHX36 stable KO cells **(Figure 4C)**. These results suggest that loss of DHX36 selectively acts to reduce C9RAN translation in a repeat length dependent manner.

### The effect of DHX36 overexpression on C9RAN DPRs expression

Since depletion of DHX36 results in a significant decrease in C9RAN translation, we wondered if overexpression of DHX36 enhances C9RAN. To address this, we expressed either a DHX36 WT or a DHX36-E335A mutant which lacks the helicase activity required to unwind G4 structures in parental HeLa cells. To ascertain the impact of DHX36 on translation in particular, we conducted studies using transfected *in vitro* transcribed C9RAN reporter mRNAs. In HeLa cells, overexpression of DHX36 significantly increased C9RAN from transfected reporter RNAs in all three sense reading frames. This effect was specific to DHX36 WT, as DHX36-E335A has no effect on C9RAN DPRs production when normalized to the FFLuc mRNA reporters and western blot analysis confirmed this relationship **(Figure 5).** These data suggest that DHX36 acts post-transcriptionally to enhance RAN translation.

**Figure 5:**
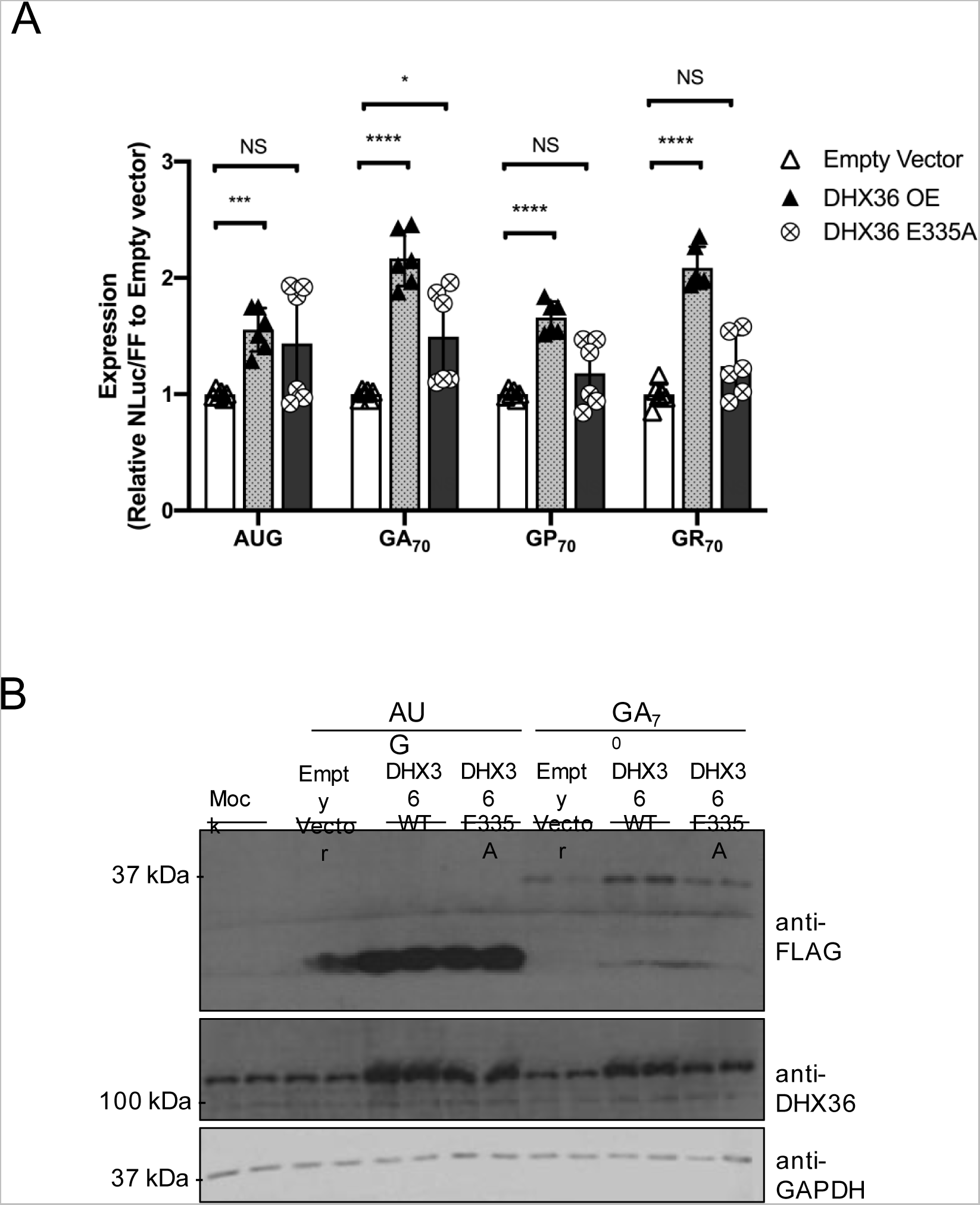
DHX36 overexpression enhances C9RAN reporter expression from G_4_C_2_ repeat RNA. (A) Relative expression of AUG and C9-RAN translation when co- transfecting reporter RNA and overexpression of empty vector, WT or E335A DHX36 DNA plasmids in HeLa cells. Data are represented as mean ±SD, n=9. Two-tailed Student’s t-test with Bonferroni and Welch’s correction, \**P* < 0.05; \*\*\**P* < 0.001; \*\*\*\**P* < 0.0001. (B) Western blots analysis of co-transfected AUG and C9-RAN luciferase reporters in RNA and empty vector, DHX36 WT or DHX36 E335A DNA plasmids in HeLa cells. GAPDH was blotted as internal control.

### Knockdown of DHX36 prevents stress dependent upregulation of RAN translation

Activation of the integrated stress response (ISR), which triggers phosphorylation of eIF2α and formation of stress granules (SGs), suppresses global protein translation initiation by impairing ternary complex recycling^71–75^. Paradoxically, ISR activation enhances RAN translation from both CGG and G_4_C_2_ repeats, and repeat expression in isolation can trigger SG formation^29, 30, 32, 37^. Loss of DHX36 induces spontaneous stress granule formation, suggesting that DHX36 may play a role in G4 structure-induced cellular stress^53^. We therefore wondered what impact loss of DHX36 would have on regulation of RAN translation in the setting of ISR activation. We co-transfected C9- NLuc and FFLuc into DHX36 control or DHX36 KD HeLa cells and then treated them with 2 μm of the ER stress inducer Thapsigargin (Tg) for 5 hrs. Tg treatment decreased expression of FFLuc in both Ctrl cells and in DHX36 KD cells, which is consistent with appropriate activation of the ISR in these cells **(Figure 6A, right)**. Consistent with prior studies^29, 37^, Tg treatment in DHX36 control cells elevated C9RAN reporter levels compared to DMSO treatment. However, depletion of DHX36 precluded this upregulation in C9RAN by Tg **(Figure 6A).** Similar findings were also observed by immunoblot in studies where we co-transfected the C9RAN reporters and GFP as a control for transfection and AUG initiated translation (**Figure 6B**). These data suggest that DHX36 may play a role in regulating the stress induction of RAN translation induced by the ISR.

**Figure 6:**
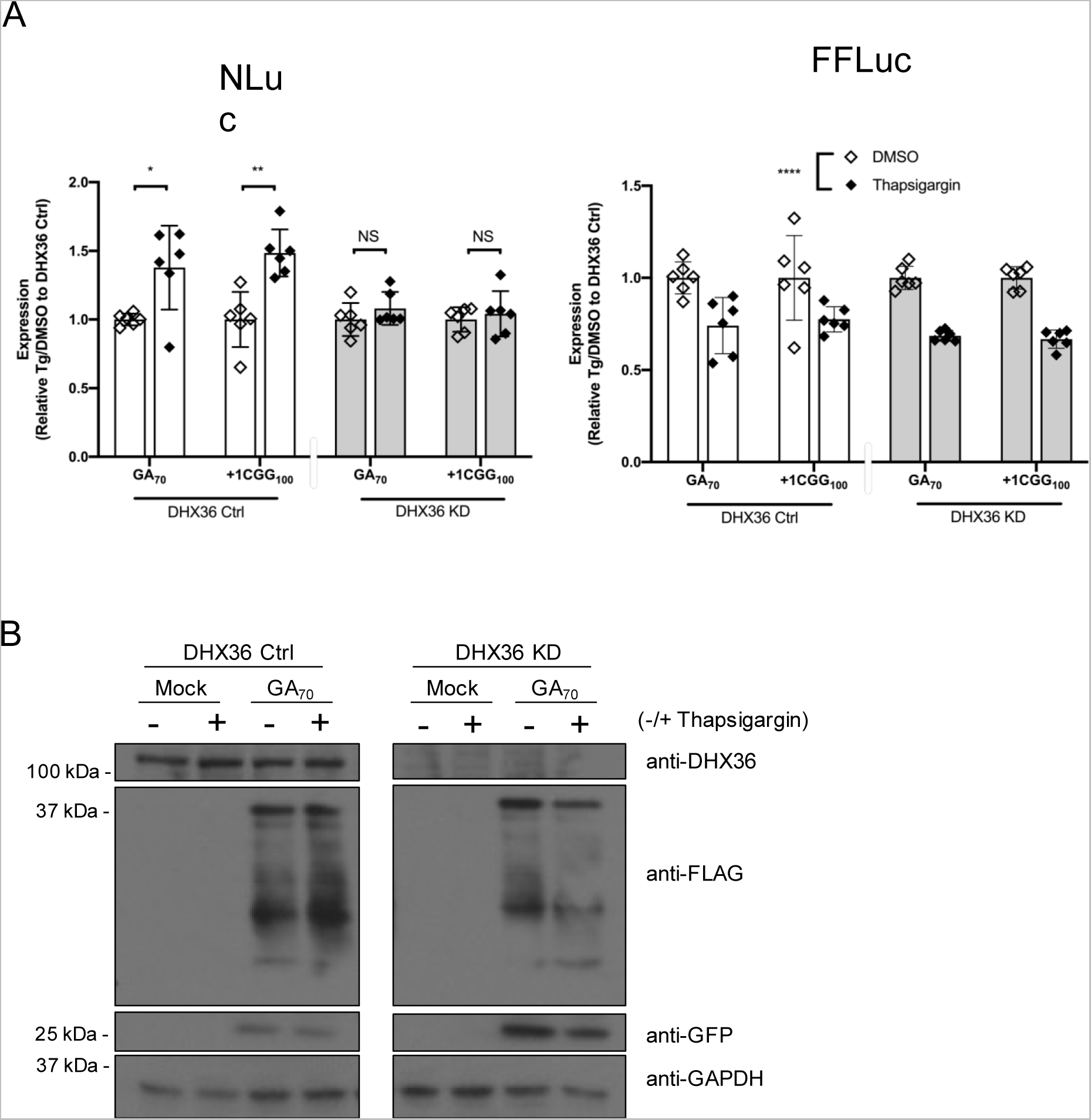
Knockdown of DHX36 prevents stress dependent upregulation of C9RAN reporter expression. (A) Relative expression of RAN translation in G_4_C_2_ and CGG repeat treated with 2 µM thapsigargin or DMSO in Ctrl and DHX36 KD HeLa cells. NLuc (left) and FF (right) signal were represented as ratio of Thapsigargin treated cells to DMSO treated cells and compared between Ctrl and DHX36 KD HeLa cells. (B) Immunoblots of G_4_C_2_ and CGG RAN luciferase reporters in Ctrl and DHX36 KD HeLa cells treated with 2 µM Thapsigargin. GFP was blotted as a transfection control and GAPDH was blotted as internal control. For panel A, data are represented as mean ±SD, n=6. two-way ANOVA was performed to discern effect of thapsigargin treatment across cell types. Two-tailed Student’s t-test with Bonferroni and Welch’s correction were performed to assess differences between individual groups. \**P* < 0.05; \*\**P* < 0.01; \*\*\*\**P* < 0.0001.

## DISCUSSION

DNA and RNA G-quadruplex (G4) structures strongly influence both gene transcription and mRNA stability, localization and translation. Moreover, G4 structures are implicated in a number of human disorders, including C9 FTD/ALS^76, 77^. Here we find that a major human G4 helicase, DHX36, enhances C9RAN translation from expanded G_4_C_2_ repeat reporter RNAs in human cells. These effects on RAN translation require DHX36 helicase activity on G4 RNA. DHX36 is also required for efficient C9RAN and CGG RAN translation as knockdown or knockout of DHX36 in human cells suppressed RAN translation from both G_4_C_2_ and CGG repeats. We also observe a robust suppression of *in vitro* C9RAN translation in DHX36 KO cell derived lysates. Overall, these data are consistent with a model whereby DHX36 binds to and unwinds GC rich repeat RNA structures and enhances their non-AUG initiated translation with a potential secondary role in enhancing repeat transcription **(Figure 7)**.

**Figure 7:**
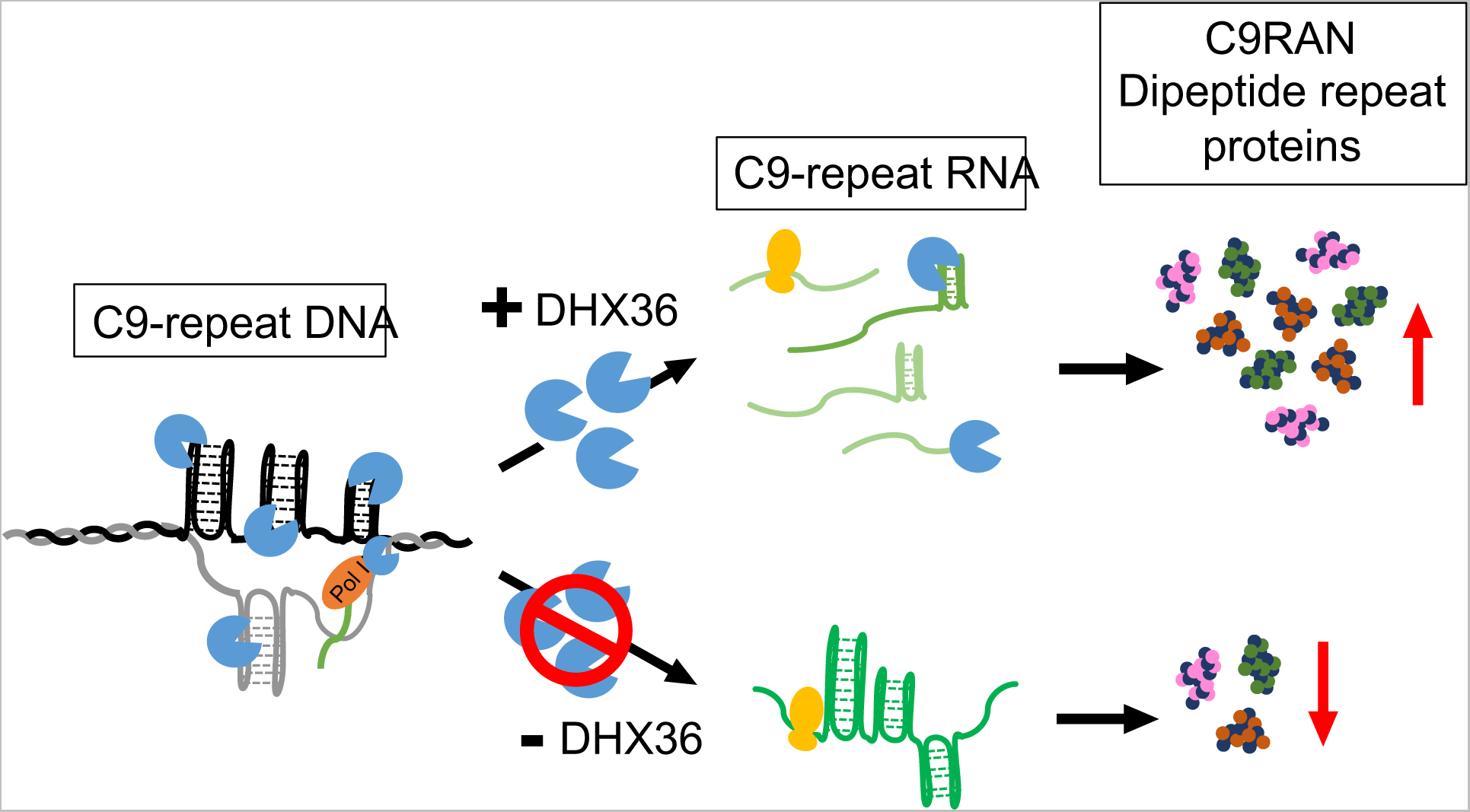
Model of DHX36 modulation of C9 RAN translation. DHX36 binds to G-quadruplex DNA and RNA structures. DHX36 aids transcription through large stretches of G_4_C_2_ repeat RNA *in vitro* but its effects in human cells with large repeats are unclear. Depletion of DHX36 decreases RAN translation from both CGG and C9-repeat reporters- both of which are capable of forming G-quadruplex structures- while increased DHX36 expression enhances C9RAN translation. These findings suggest a direct role for DHX36 in RAN translation of GC rich repeats.

The observed effects on C9RAN reporter generation are largely post-transcriptional. While we do observe a stimulatory effect of DHX36 on T7 polymerase transcription from a C9-repeat *in vitro* **(Figure 1D-E)**, we do not observe changes in C9-repeat RNA levels in cells following DHX36 KD or overexpression **(Figure 2G, Supplemental Figure 5).** We also show that DHX36 directly binds to C9-repeat DNA with a binding affinity of ∼10- 100 less than previously reported for pure G4 DNAs^40, 44, 49^ **(Figure 1A-C)**. The relatively low affinity of DHX36 for C9-repeat G4 DNA might in part explain the lack of a robust effect on C9-repeat transcript levels in cells following DHX36 KD or overexpression. In addition, T7 polymerase *in vitro* is prone to early transcription termination^78^ and as such may be less efficient than RNA polymerase complexes at resolving RNA structures and generating complete transcripts in cells. Future work using patient-derived cells harboring greater repeat lengths (which DHX36 may have greater affinity for) will be necessary to more fully characterize the potential for DHX36 to modulate C9-repeat transcription in patients.

DHX36 binds to C9-repeat RNA in a G4 specific manner both *in vitro* and in studies using human cell and mouse spinal cord lysates^6, 51, 79^. Depletion of DHX36 decreases C9RAN translation and this decrease occurs across all reading frames and is dependent on the length of the repeats **(Figures 2-4)**. Similar results are seen for CGG repeats capable of supporting RAN translation and folding into G4 structures. This suggests that RAN translation initiation or elongation could be significantly influenced by both RNA binding protein recognition and resolution of repeat RNA secondary structures. This idea is supported by the finding that a helicase dead form of DHX36 failed to influence RAN translation of C9-repeat RNAs. It is also consistent with prior studies implicating RNA helicases such as DDX3X and the eIF4A helicase cofactors eIF4B and eIF4H as modifiers of RAN translation at both CGG and G_4_C_2_ repeats^33, 36^.

In addition, depletion of DHX36 precluded the augmentation of RAN translation typically observed in response to stress **(Figure 6)**^29, 30, 80^. DHX36 is a component of stress granules and plays an active role in regulating the cellular stress response^81–83^. Indeed, KD or KO of DHX36 is sufficient to trigger stress granule formation without application of an exogenous stressor^53^. How exactly loss of DHX36 precludes this upregulation is not clear. ISR activation augments RAN translation at least in part by lowering initiation codon fidelity requirements^29, 31^. If DHX36 is specifically influencing elongation through the repeat, then its depletion may slow translation due to ribosomal stalling within the repeats despite continued enhanced initiation. Alternatively, G_4_C_2_ repeats also support a 5’ M^7^G cap independent “IRES-like” RAN initiation mechanism that is enhanced by ISR activation^30, 84–86^. DHX36 could play an active role in generating this structure and allowing for internal ribosome entry. A deeper understanding of RNA structure/function relationships as they apply to RAN translation will be needed to determine which of these mechanisms (or both) is likely to explain how DHX36 loss impacts RAN translation at both CGG and G_4_C_2_ repeats.

This study has some limitations. It largely relies on reporter assays using transiently transfected plasmids or *in vitro* transcribed linear RNA. RAN translation from the endogenous locus of C9 might involve very large RNA from longer repeats and the exact nature of the RAN translated transcripts in C9 patient neurons remains unclear. In particular, endogenous RNAs may form a combination of dynamic secondary structures including hairpins and G4s, which complicate the potential effects on both RNA mediated toxicity and RAN translation. Further studies using endogenous systems such as C9 FTD/ALS patient iPSC derived neurons and rodent that harbor larger repeats will be needed to confirm the roles of DHX36 in endogenous repeat transcription, RAN translation, and toxicity derived from the endogenous repeat.

In sum, this study provides evidence that DHX36 can influence RAN translation of G_4_C_2_ repeats both basally and in response to stress pathways. These studies suggest that control of G4 formation at the DNA and RNA levels and modulation of G4 resolving helicases such as DHX36 are candidate therapeutic strategies and targets for G rich repeat-associated neurological diseases.

## METHODS

### Electrophoretic mobility shift assays

62.5 pg of 5’- IRDye 800 labeled*C9* or scrambled oligonucleotides **(Table 1)** were synthesized (Integrated DNA Technologies, Inc.) were heated in nuclease-free water in the presence of KCl (100 mM) starting at 98°C and decreasing 10°C every two min, ending at 28°C. As a control, 62.5 pg of *C9* oligonucleotides were heated and cooled as above in the absence of KCl, 3.75 pg of each oligonucleotide were incubated at 37°C for 30 min in binding buffer (75 mM EDTA, 40.6 mM Tris-Acetate pH 7.8, 40.6 mM NaCl, 1.21 mM MgCl2, 8.1% glycerol, 0.0065% lactalbumin, 2-mercaptoethanol, 0.16x protease inhibitor cocktail (Roche), 0.12 mg/mL leupeptin) with varying concentrations of rDHX36 (7.4, 4.5, 1.7, 1.1, or 0 nM). Additional volumes of buffer were added to the 4.5, 1.7, 1.1, or 0 nM reactions so as to have equal buffer concentrations in all reactions. Glycerol (16% final) was added to the samples, and 3 pg of DNA were loaded per well onto a 10% non-denaturing PAGE. The samples were electrophoresed for 5 hrs at 120 V in the dark. Each EMSA was performed in triplicate and analyzed using a Li-Cor Odyssey imager. Percent bound was determined by densitometry measurements in ImageStudio using the following equation: percent bound = (bound DNA / total DNA) x 100. Triplicate values were averaged and plotted with GraphPad Prism using a non-linear regression (curve fit) function.

**Table 1.**
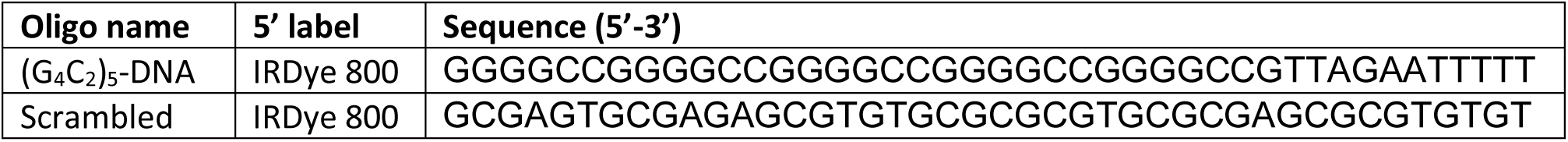
DNA oligonucleotides used in EMSA studies.

### *In vitro* transcription assay

A HiScribe™ T7 Quick High Yield RNA Synthesis Kit (New England Bio Labs, Cat.# E2050S) was used with 1 µg of a GA_70_ or AUG-NLuc linearized plasmid as template^29^. The reactions were transcribed for 15 hrs at 37°C in the presence of absence of DHX36 (0 or 6 nM). An additional volume of DHX36 storage buffer was added to reactions without DHX36 so as to have equal buffer concentrations in all reactions. The resulting transcripts were treated with DNase for 20 min at room temperature and then isolated using Micro Bio-Spin™ 6 Columns (BioRad, Cat.# 732-6221). 0.75 µg of RNA transcripts were mixed with 2X formamide buffer (95% deionized formamide, 0.025% bromophenol blue, 5 mM EDTA) and heated at 95°C for 5 min. The samples were resolved in a 7.5% denaturing-8M urea PAGE for 20 min at 2 W and then 45 minutes at 20 W. The gels were subsequently soaked in SYBR Gold nucleic acid stains (ThermoFisher, Cat.# S11494) for 20 min at RT and imaged using a BIO-Rad Gel Docs™ XR+ and quantified using densitometry software. Densitometry values for the top third of the gel were divided by total value for each lane. Values for total densitometry readings for each lane were also taken.

### RNA synthesis

pcDNA3.1(+)/NLuc-3xF and pcDNA3.1(+)/FF were linearized by PspOMI and XbaI restriction enzymes respectively and recovered using DNA Clean and Concentrator-25 kits (Zymo Research, Cat# D4033). m^7^G-capped and poly-adenylated RNAs were transcribed in vitro from these plasmids using HiScribe T7 ARCA mRNA Kit, with polyA tailing (NEB, Cat# E2065S) following the manufacturer’s instructions and recovered using RNA Clean and Concentrator-25 kits (Zymo Research, Cat# R1017). The integrity and size of all transcribed RNAs were confirmed by denaturing formaldehyde and formamide agarose gel electrophoresis.

### Cell culture, transfection, qRT-PCR, and drug treatment

Jurkat T1-28/11 and HeLa 15/25 cells were cultured and passaged at 37°C, 5% CO_2_. Jurkat T1-28/11 cells were maintained in Roswell Park Memorial Institute (RPMI) 1640 Medium supplemented with 10% fetal bovine serum (FBS). HeLa 15/25 cells were maintained in Dulbecco’s modified Eagle Medium (DMEM) supplemented with 10% FBS, 1% non-essential amino acids, 3 μg/ml blasticidine, and 250 ug/ml zeocin. To induce DHX36 knockdown, HeLa 15 and HeLa 25 were both treated with daily changed media contained 1 ug/ml doxycycline (doxy.) for 4 days.

For transfection and luciferase assay in Jurkat cells, cells were plated in 24-well plates at 6x10^5^ cells/well in 500 μl media. Reverse transfection by using TransIT-Jurkat transfection reagent (Mirus, Cat.# MIR 2124) was done after cells plating. Cells were co-transfected with 250 ng/well of pcDNA(+)-NLuc-3xFLAG plasmids developed from Kearse et al. and Green et al. and 250 ng/well of pGL4.13 firefly luciferase plasmid as transfection control^29, 69^. Mixed plasmids and reagents were added drop-wise in cultured cells after 30 min incubation at room temperature, then the plate was gently shaken for 1 min. Luciferase assays were performed 48 hrs after plasmid transfection. Cells from each well were collected in microcentrifuge tube and media was removed after 400 rpm centrifuge for 5 min. Then cells from each tube were lysed with 60 μL of Glo Lysis Buffer 1X (Promega, Cat.# E2661) and were vortexed for 5 sec. In opaque white 96-well plates, from 60 μL of cell lysate, 25 μL of cell lysate was distributed to mix with 25 μL of Nano-Glo Luciferase Assay System (Promega, Cat.# N1120), and another 25 μL of cell lysate was mixed with 25 μL of ONE-Glo Luciferase Assay System (Promega, Cat.# E6130). The plate was placed on a shaker for 5 min in the dark. Luciferase activity in each well was obtained by luminescence measurements. All reagents and experiments are presented at room temperature.

For transfection and luciferase assay in HeLa cells, cells were plated in 96-well plates at 2.5x10^5^ cells/well in 100 μl media. 24 hrs after plating, cells were co-transfected with 50 ng/well of pcDNA(+)-NLuc-3xFLAG plasmids, and 50 ng/well of pGL4.13 firefly luciferase plasmid as transfection control. Transfection was done by adding Viafect transfection reagent (Promega, Cat# E4981) with mixed plasmids drop-wise in cultured cells after 10 min incubation at room temperature then gently shaking the plate for 1 min. Plasmid DNA and C9-repeat RNA co-transfection were done by forward transfection of published DNA plasmid expressing empty vector, WT, or E335A DHX36^41^ in HeLa cells seeded at 2.5x10^5^ cells/well in 100 μl media. After 24 hrs, *in vitro* synthesized C9-RNA and pcDNA-FF RNA were co-transfected at 50 ng/well each into the well by Viafect transfection reagent (Promega, Cat# E4981) as described above. Following luciferase assays were performed 24 hrs after C9-repeat DNA plasmids or RNA transfection, as described by Kearse et al^69^.

For qRT-PCR assays, after HeLa cells were plated and transfected as described above, experiments were performed as described by Linsalata et al^33^.

For C9RAN reporter luciferase analysis following stress activation, after 4 days of doxy. treatment in HeLa 15/25 cells, cells were seeded and transfect for 19 hrs then followed by 5 hrs treatment of 2 μM Thapsigargin (Tg).

### Immunoblot and antibodies

In a 12-well plate, HeLa 15/25 cells were rinsed with 500 μL cold 1X PBS twice then lysed in 300 μL RIPA buffer with protease inhibitor (120 μL for 24-well plates) for 30 min at a 4°C shaker. Lysates were homogenized by passing through a 28G syringe 8 times, mixed with 6X sample buffer with a final of 2% B-ME, denatured at 95°C for 10 min, and stored at -20 °C. Protein samples were standardized by BCA assay for equal total protein loading. 20 μL of equal total protein sample was loaded in each well of a 10% SDS-PAGE. All primary antibodies applied for Western Blot were used at 1:1000 in 5% non-fat dairy milk (wt/vol) and 0.1% Tween-20 (vol/vol) in TBS except anti-puromycin at 1:5000. Monoclonal mouse anti-DHX36 antibody was generated at 2.57 μg/μl^41^, monoclonal mouse anti-FLAG antibody was from Sigma (clone M2, Cat#F1804), mouse anti-GFP was from Roche (Cat# 11814460001), monoclonal mouse anti-GAPDH was from Santa Cruz Biotechnology (clone 6C5, Cat# sc-32233), mouse anti-puromycin 12D10 was from Millipore (Cat# MABE434).

### Jurkat cell line generation

The DHX36 gRNAs (T1, T2, and T10) were designed to target exonic regions of DHX36/G4R1 (Gene ID: 170506) in order to disrupt all the gene products (**Supplemental Figure 3**). The gRNA T1 (AAGTACGATATGACTAACAC) was evaluated to be the most effective by nucleotide mismatching assay in the cell pool examination (31.2% cleavage efficiency) and was utilized for generation of single cell clones. Cleavage efficiency was determined by sequencing trace analysis with the online tool TIDE (https://tide-calculator.nki.nl/). Clones were identified and confirmed using Sanger Sequencing of PCR and RT-PCR productions (**Supplemental Figure 3**) and western blot analysis (**Figure 3, A**)

### In vitro translation assays

Preparation of cell lysate, hypertonic lysis buffer, and translation buffer were followed by Linsalata et al ^33^. Jurkat cells were centrifuged, and rinsed 3 times with PBS (pH 7.4). Hypotonic lysis buffer that contained 10 mM HEPES-KOH (pH 7.6), 10 mM KOAc, 0.5 mM Mg2OAc, 5 mM DTT, and EDTA-free protease inhibitor was added to cells pellet on ice to swollen the cells for 30 minutes. Then cells were mechanically lysed by 20 strokes in a 27G syringe and followed by another 30 minutes of incubation on ice. The supernatant from the cell lysate was collected by centrifuging the cell lysate at 10,000 g for 10 min at 4°C and further diluted in lysis buffer to 8.0 ug/ul using a modified Bradford protein quantification assay (Bio-Rad), flash-frozen in liquid N_2_, and stored at 80°C. The lysates were added to the translation buffer that with final concentrations of 20 mM HEPES-KOH (pH 7.6), 44 mM KOAc, 2.2 mM Mg2AOc, 2 mM DTT, 20 mM creatine phosphate (Roche), 0.1 ug/ul creatine kinase (Roche), 0.1 mM spermidine, and on average 0.1 mM of each amino acid.

For in vitro translation assays, 40 fmol of RNA was added to lysate which protein concentration at 8 ug/ul of 10 ul per reaction. After incubation at 30°C for 120 min, 25 ul room temperature Glo Lysis Buffer (Promega) was added to the reaction to incubate for 5 min at room temperature. With the Nano-Glo Dual-luciferase system, 25 ul of this mixture was added 25 ul of the ONE-Glo™ EX reagent following 25 ul of NanoDLR Stop & Glo reagent (Promega). All mixtures were incubated in opaque white 96-well plates on a rocking shaker in the dark for 5 min before quantifying the luminescence.

### Measuring protein synthesis by puromycin incorporation

Nascent Global translation was monitored by the surface sensing of translation (SUnSET) method^58^. After seeding HeLa cells as described above in a 24-well plate format, 48 hrs later, cells were incubated with fresh media containing 10 μg/ml of puromycin for 10 min at room temperature. Cells were then placed on ice and washed with ice-cold PBS, prior to lysis in 100 μL RIPA buffer containing protease inhibitor.

### Statistical methods

Statistical analysis was performed using GraphPad Prism8. For comparison of NLuc and FFLuc reporter luciferase activity, one-way ANOVAs were performed to confirm statistical difference between control and experimental groups. Two-way ANOVA was performed to confirm the statistical difference on FFLuc signal from the treatment of Thapsigargin between different groups of control and experimental conditions. Post-hoc Student’s t tests were then performed with Bonferroni correction for multiple comparisons and Welch’s correction for unequal variance. All studies represent at least three independently replicated experiments. All bar graphs include a standard deviation error bar and each independent replicate. Exact N for each sample and analysis performed are noted in the figure legend.

## Acknowledgements

We thank everyone in the Todd lab and the Smaldino lab for thoughtful feedback and discussion on this project. Jiou Wang provided initial G_4_C_2_ repeat plasmids to the Smaldino Lab.

---

**Supplemental Figure 1:**
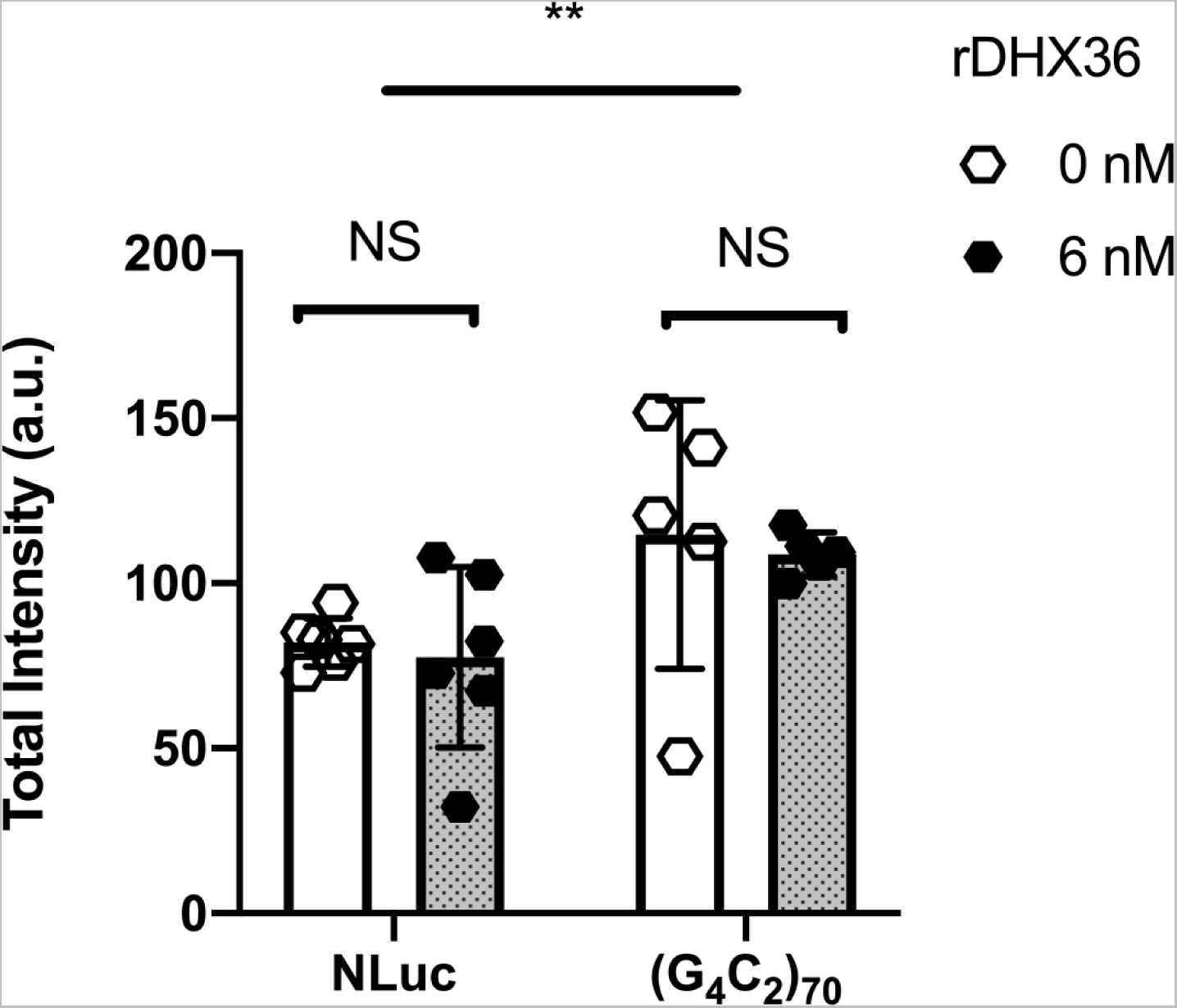
Total RNA quantification of in vitro transcription. The total density of each lane were quantified after subtraction to the background signal. Data are presented as ±SD, n=5(NLuc) - 6 (G_4_C_2_)_70_. Two-tailed pair t-test was performed to compared between with or without rDHX36, N.S. = not-significant. Two-way ANOVA was performed to compare differences between NLuc and (G_4_C_2_)_70_ experimental groups, \*\**P* < 0.005.

**Supplemental Figure 2:**
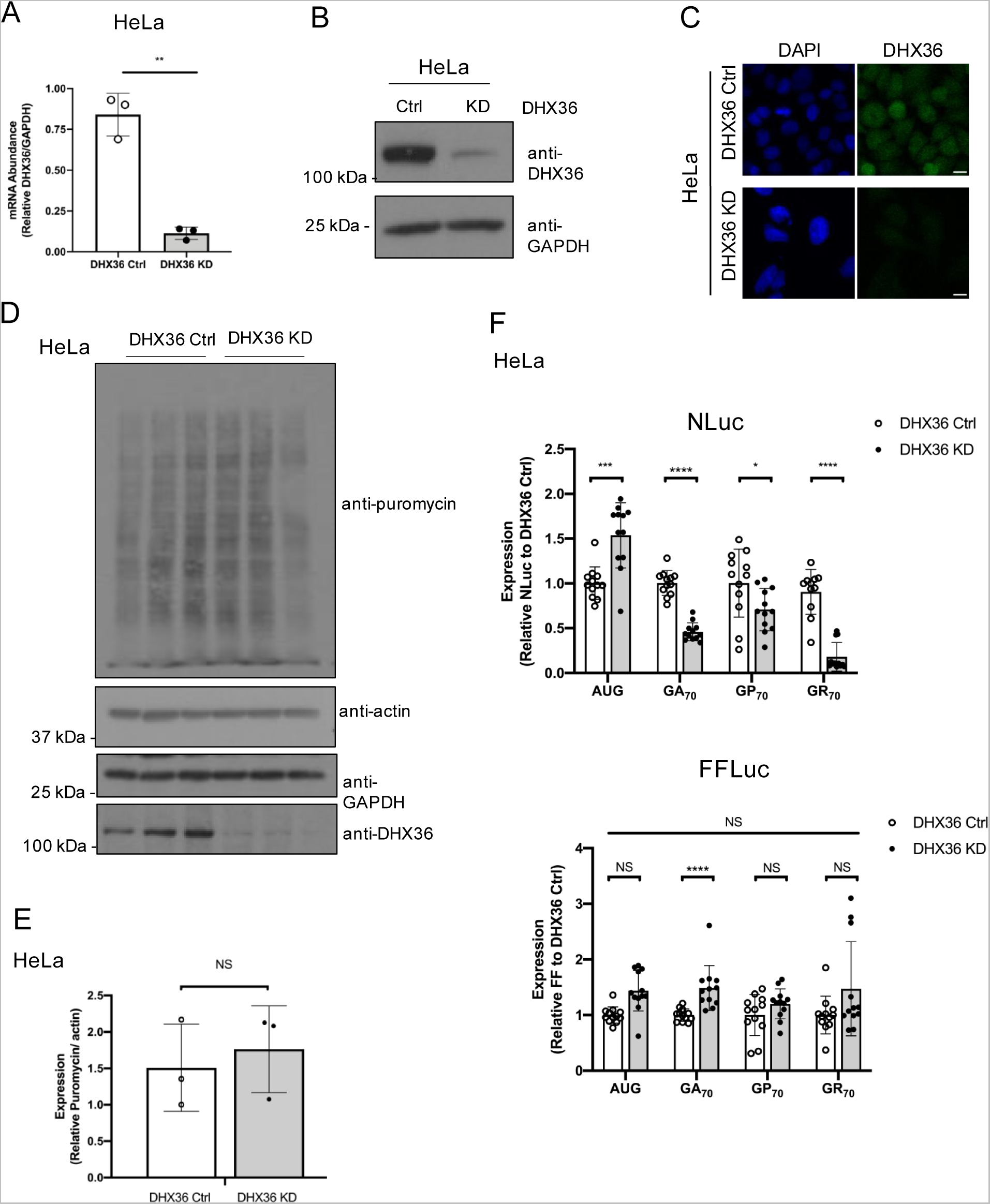
Characterization of Control and DHX36 KD in HeLa cells. Relative abundance of DHX36 mRNA in Ctrl and DHX36 KD HeLa cells as measured by qRT-PCR (normalized to GAPDH mRNA). Data are represented as mean ±SD, n=3. Two-tailed Student’s t-test with Bonferroni and Welch’s correction, \*\**P* < 0.01 (A) Western blots of DHX36 in Ctrl and DHX36 KD HeLa cells. GAPDH was blotted as internal control. (C) Immuno-fluorescent images of Ctrl and DHX36 KD HeLa cells expressing DHX36, scale bar = 10 µm. (D&E) Translation was monitored by puromycin incorporation and quantification was performed by normalizing puromycin to actin. Data are represented as mean ±SD, n=3. (F) Relative NLuc or FF signal normalized to DHX36 WT. One-way ANOVA comparing FF signal between DHX36 control and KD groups in different treatment conditions. Then two-tailed Student’s t-test with Bonferroni and Welch’s correction was further performed to test for statistical difference between AUG and C9-RAN translation groups, N.S. = not-significant; \**P* < 0.05; \*\*\**P* < 0.001; \*\*\*\**P* < 0.0001.

**Supplemental Figure 3:**
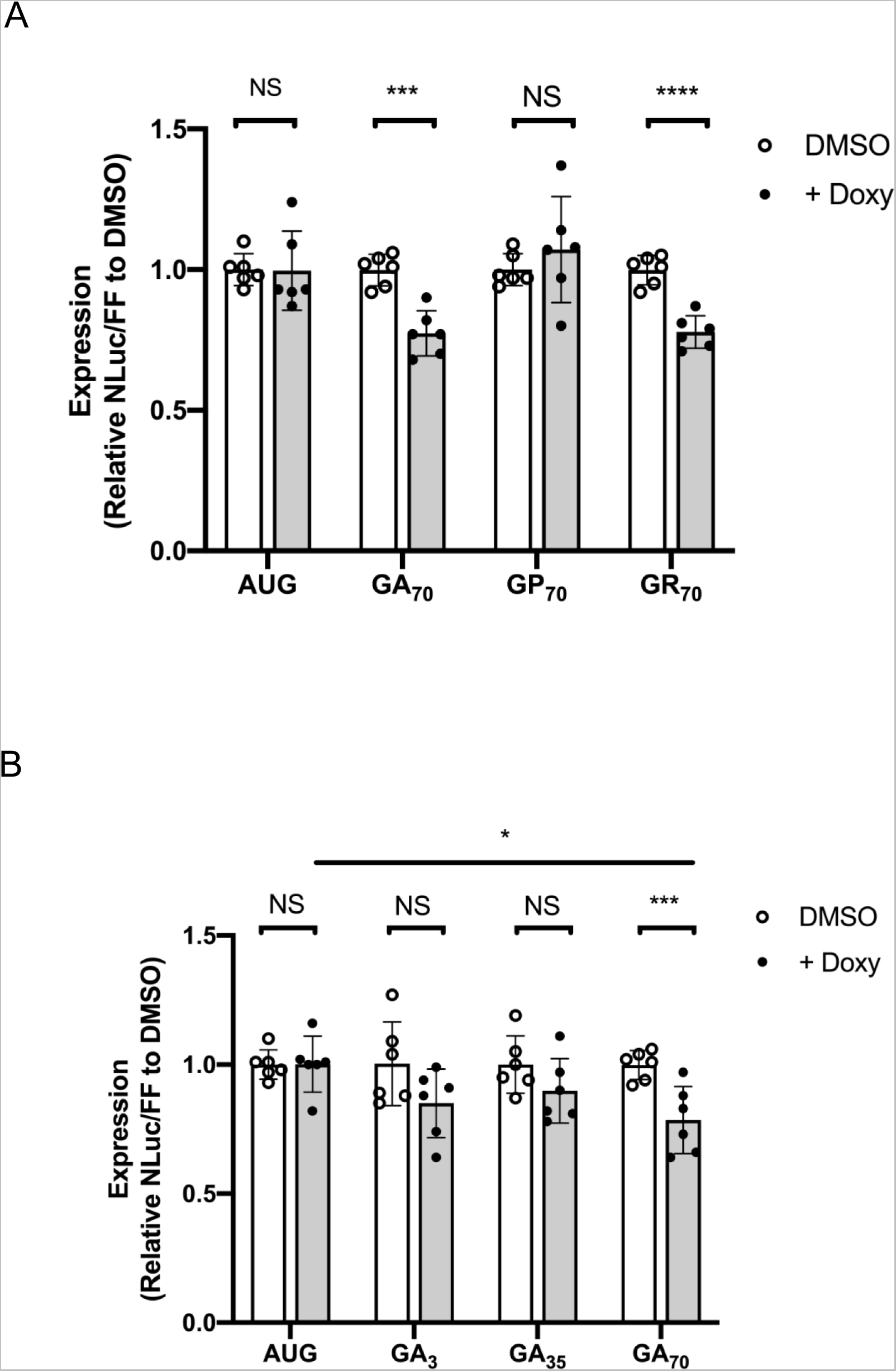
C9RAN reporter expression in DHX36 KD inducible HeLa cells. treated with DMSO or Doxycycline. Inducible DHX36 KD HeLa cells were treated with DMSO or Doxy prior plasmids transfection. (A) Relative expression of AUG and C9-RAN translation in GA (+0), GP (+1), and GR (+2) frames with 70 repeats. (B) Relative expression of AUG and C9-RAN translation from GA frames with 3, 35, and 70 repeats. NLuc signal were normalized to AUG-FF translation. Data are represented as mean ±SD, n=6. One-way ANOVA was performed to compare the statistical differences between repeat length in treatment of Doxycycline. Two-tailed Student’s t-test with Bonferroni and Welch’s correction, \**P* < 0.05; \*\*\**P* < 0.001; \*\*\*\**P* < 0.0001.

**Supplemental Figure 4:**
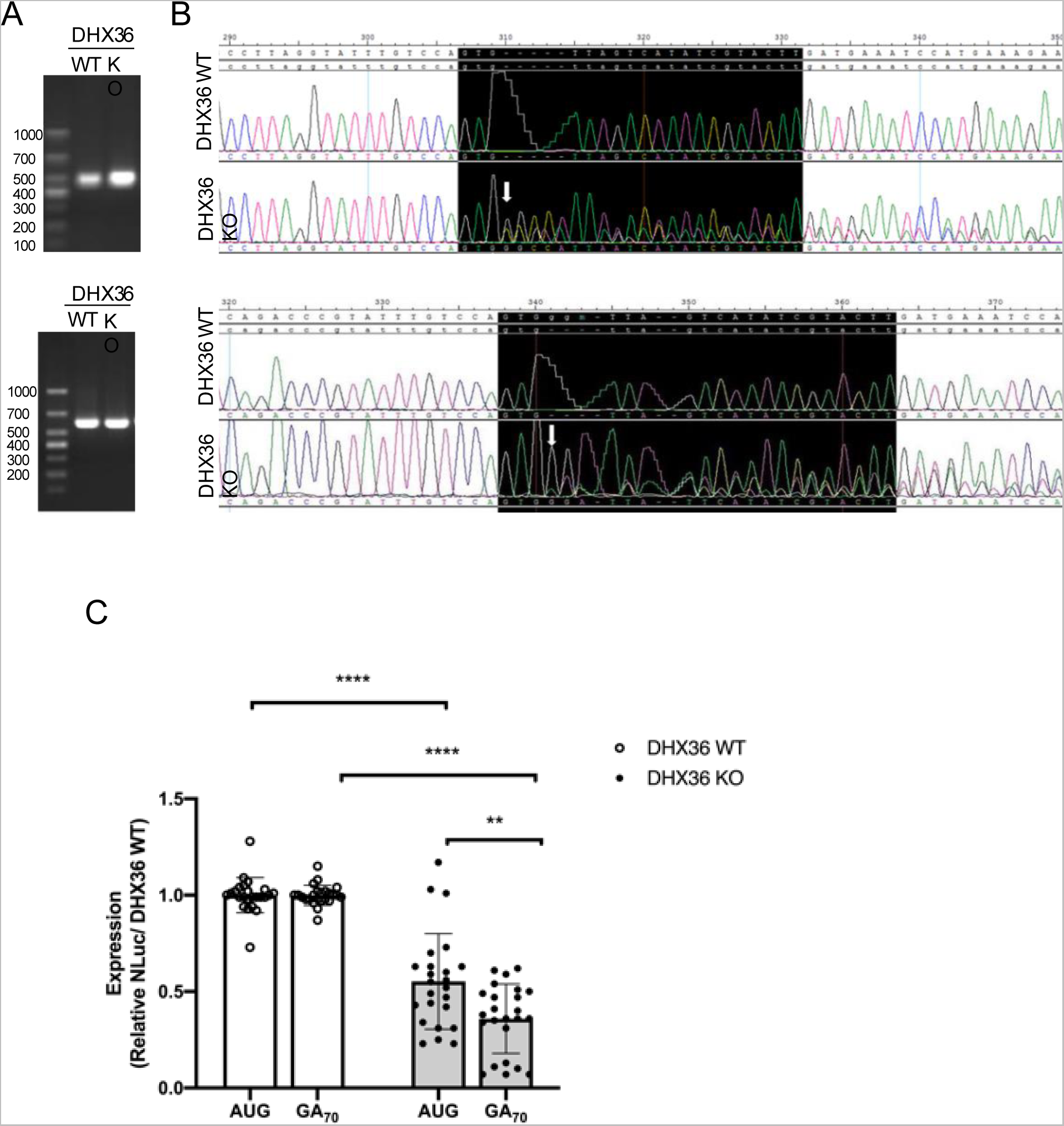
Generation of DHX36 Knockout Jurkat cell lines. (A) PCR (top) and RT-PCR products (bottom) from primers flanking the DHX36 gRNA site (B) Alignment of Sanger sequencing data from PCR (top) and RT-PCR (bottom) products. gRNA targeting site was shaded in black. Arrow indicates the site of INDEL mutation. The DHX36 WT clone was identified and confirmed as a negative KO clone with genotype of the target gene identical to the parental cell line (INDEL:0/0). The DHX36 KO clone was identified and confirmed with a 5 bp insertion on one allele and 6 bp insertion on the other allele at the targeted site (INDEL: +5/+6). (C) *In vitro* translation assay demonstrates knockout of DHX36 preferentially decreases C9-RAN translation compared to AUG-NLUC control. NLuc signal were normalized to DHX36 WT. Data are represented as mean ±SD, n=24. Two-tailed Student’s t-test with Bonferroni and Welch’s correction, \*\**P* < 0.01; \*\*\*\**P* < 0.0001.

**Supplemental Figure 5:**
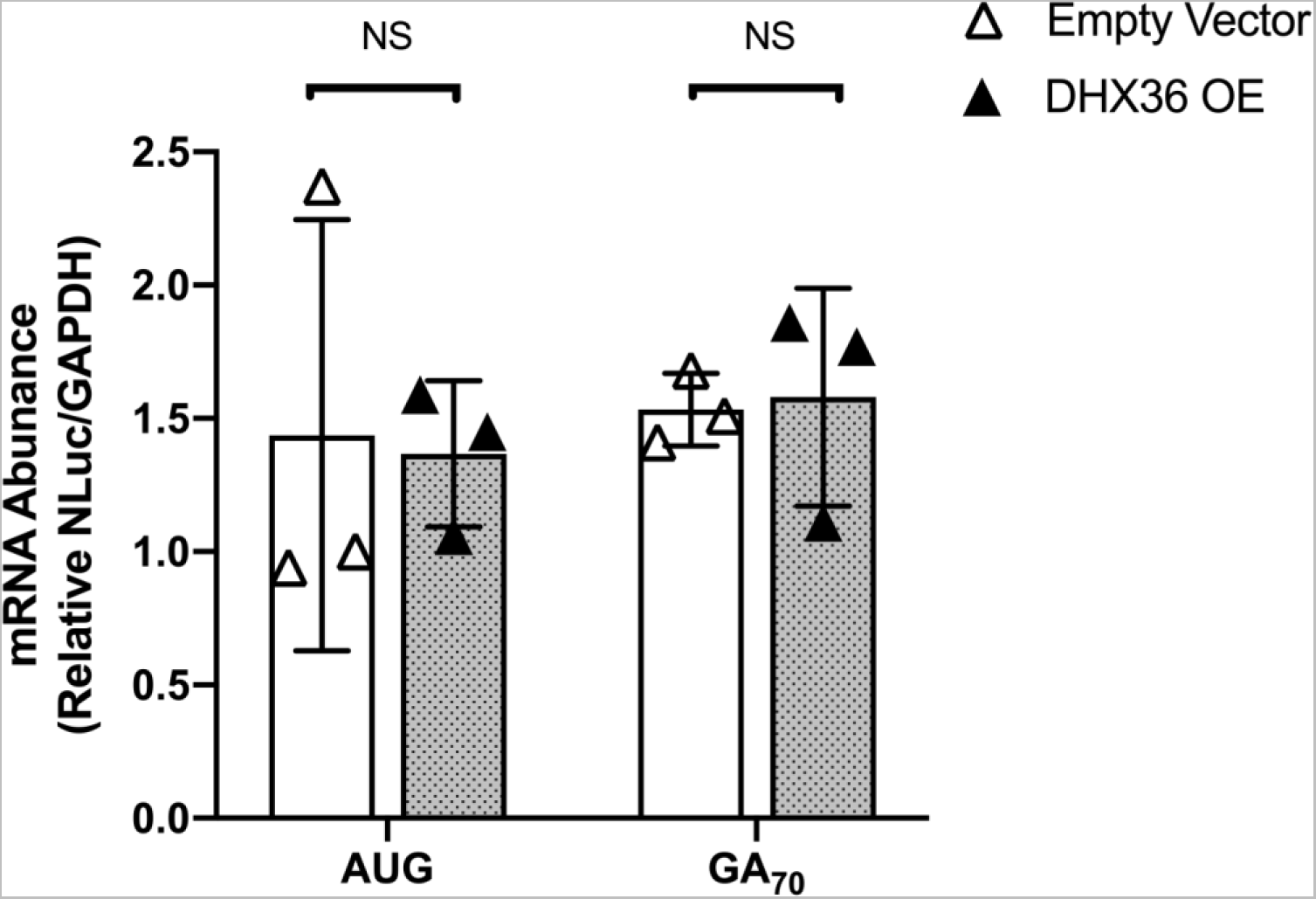
DHX36 overexpression effect on C9RAN mRNA abundance from transfected plasmids. Relative expression of AUG and C9-RAN translation when co-transfecting reporter DNA and overexpression of empty vector, DHX36 WT or DHX36 E335A DNA plasmids in HeLa cells. Data are represented as mean ±SD, n=9. Two-tailed Student’s t-test with Bonferroni and Welch’s correction, \**P* < 0.05; \*\**P* < 0.01; \*\*\**P* < 0.001. (B) qRT-PCR detecting NLuc mRNA from cells transfected with empty vector or DHX36 overexpression in HeLa cells. Data are represented as mean ±SD, n=3. Two-tailed Student’s t-test with Bonferroni and Welch’s correction, not significant.

**Supplemental Figure 6:**
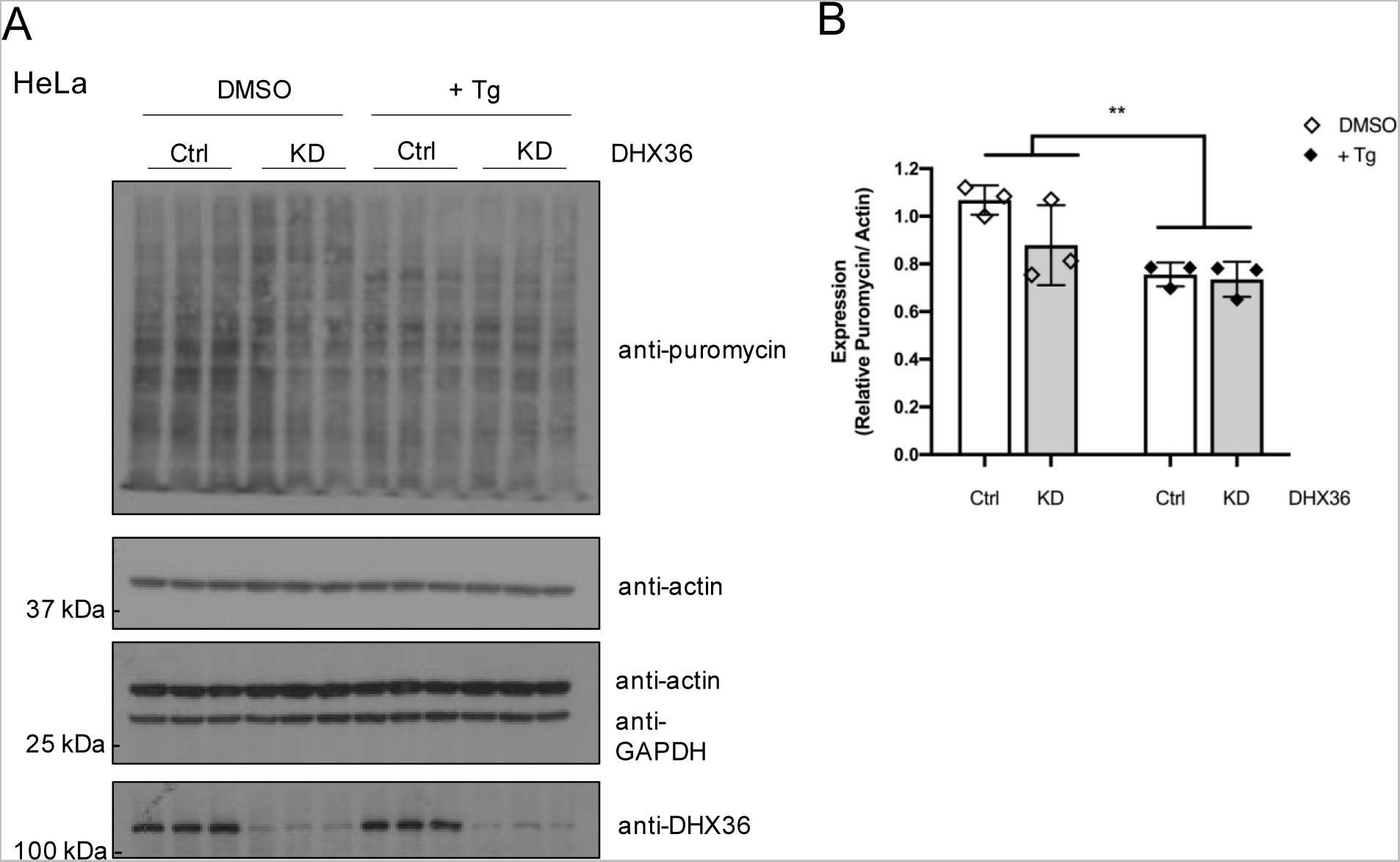
Global protein translation in DHX36 KD HeLa cells under Thapsigargin stress. (A-B) Effect on protein synthesis under Tg stress was monitored by puromycin incorporation and quantification was performed by normalizing puromycin to actin. With treatment of Tg, there is a decrease on global translation. Data are represented as mean ±SD, n=3. One-way ANOVA compared between DMSO and Tg treated groups, \*\**P* < 0.01.

**Figure 1.**
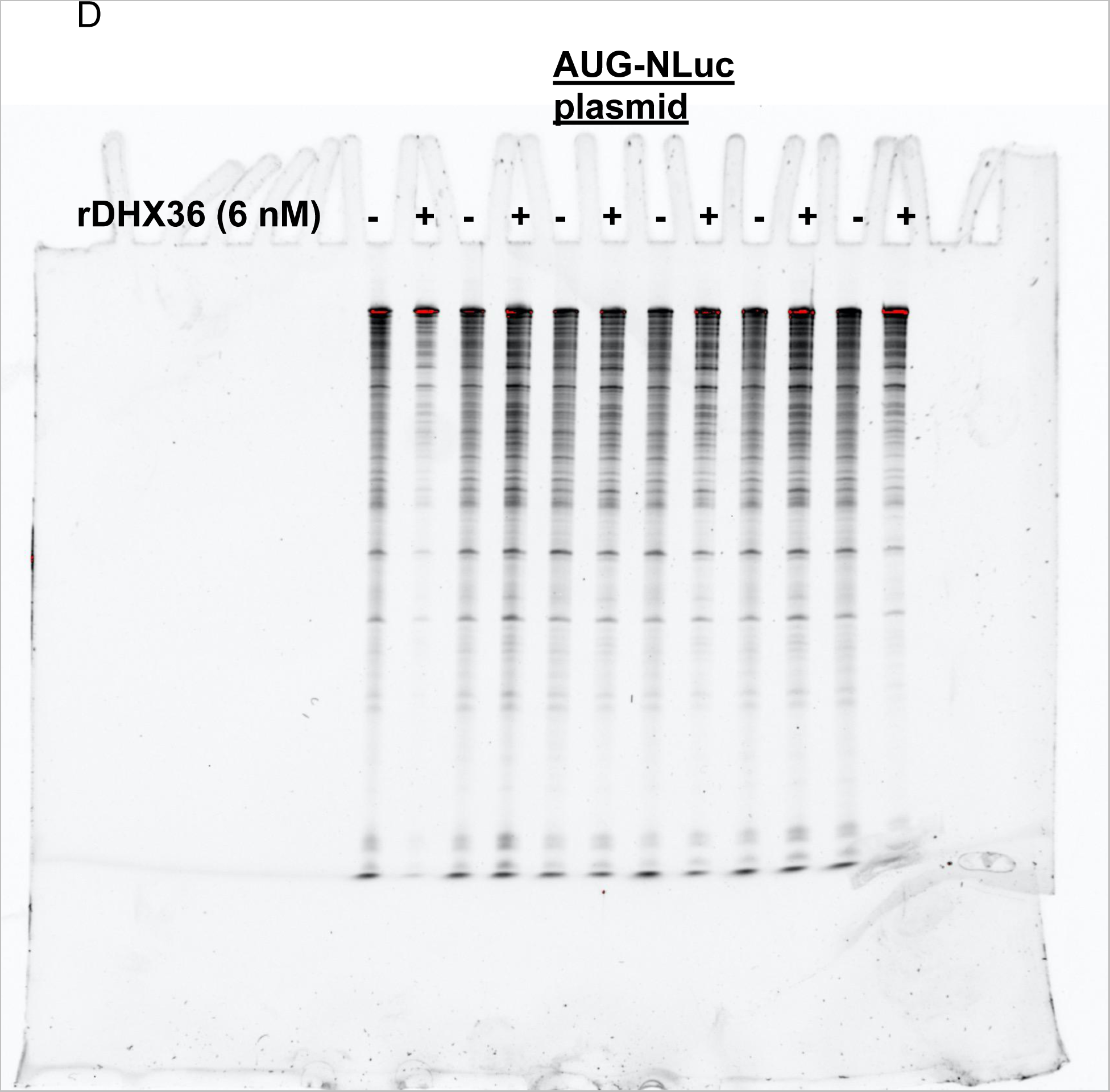
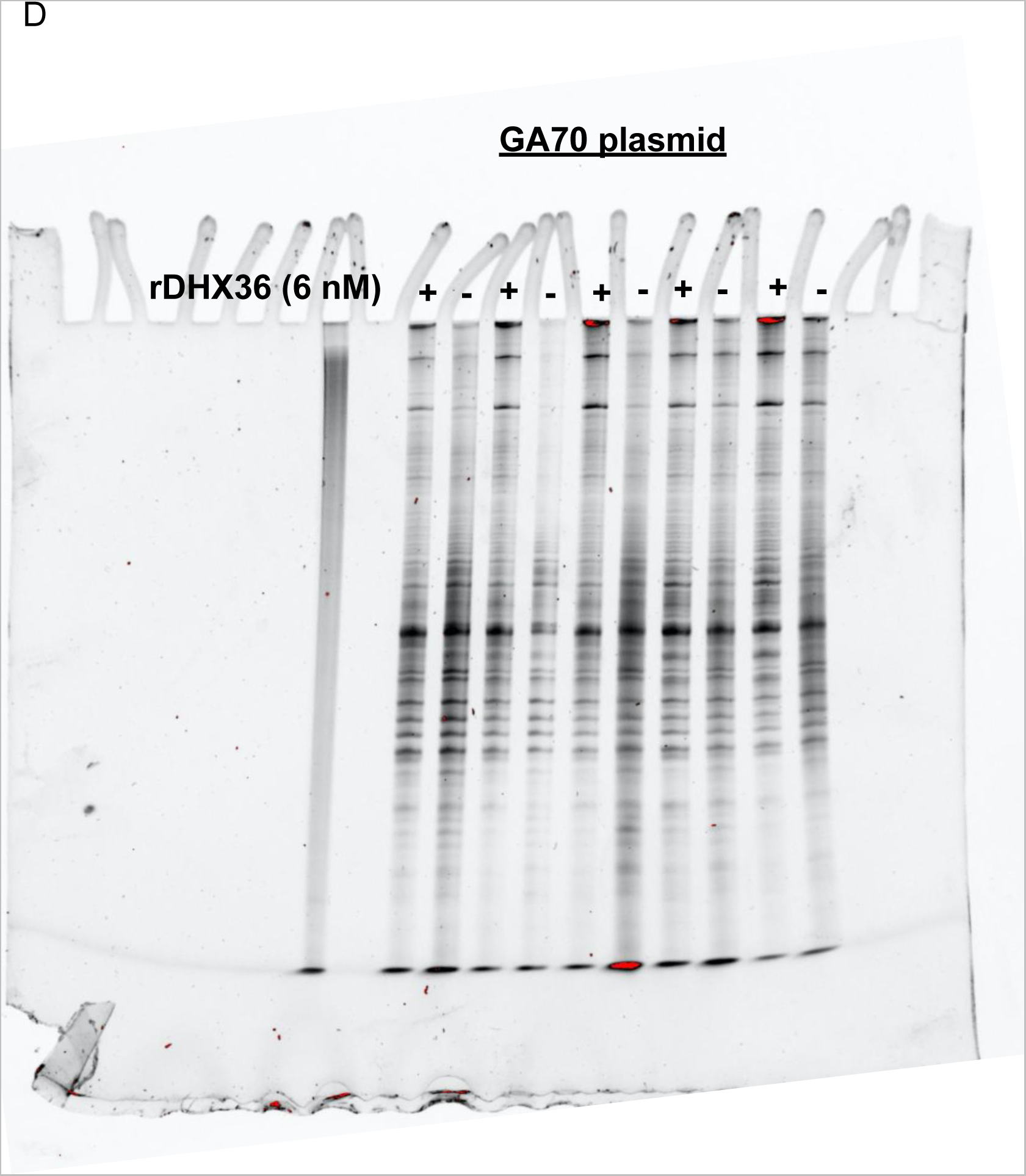

**Figure 2.**
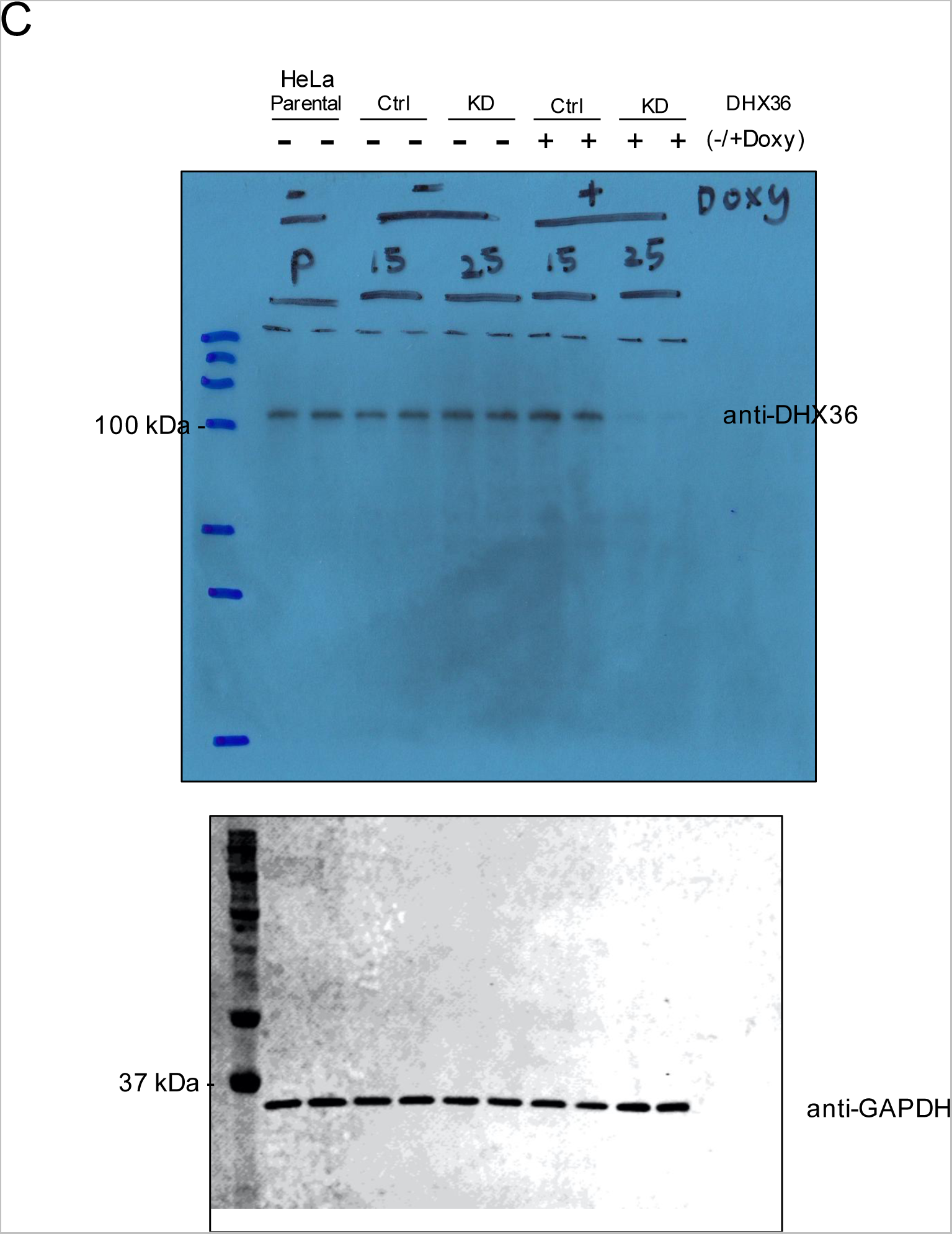
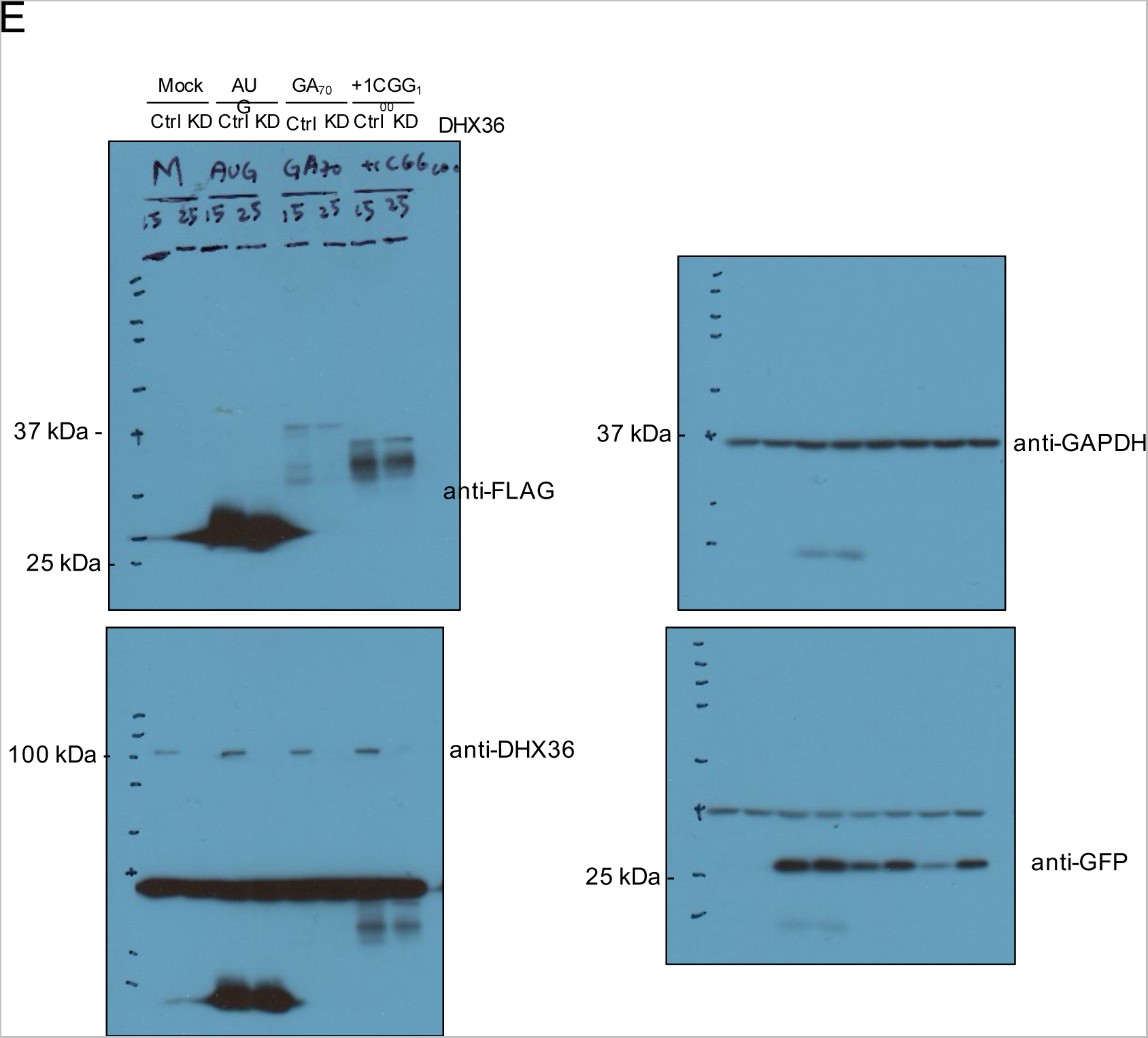

**Figure 3.**
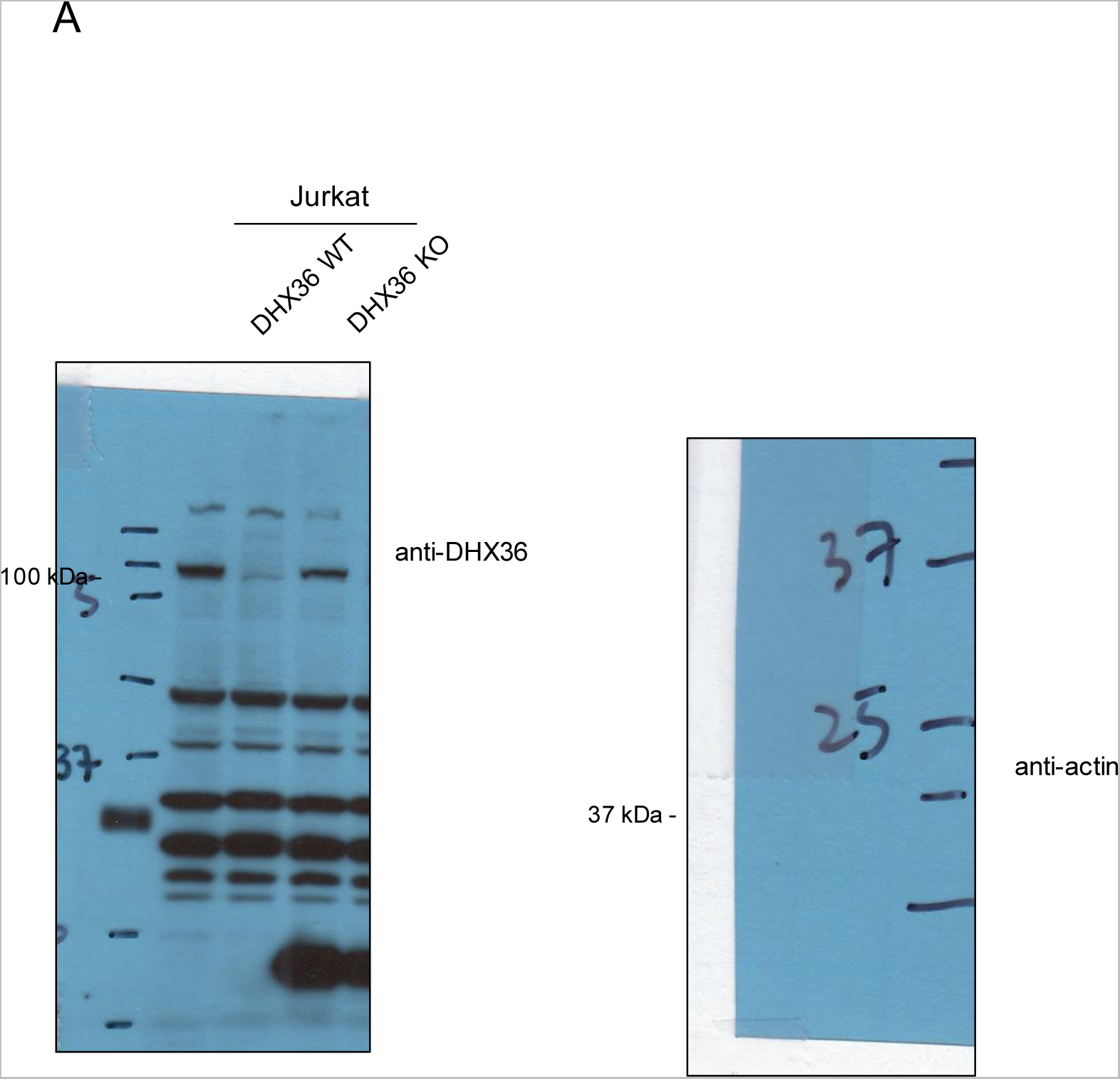

**Figure 5.**
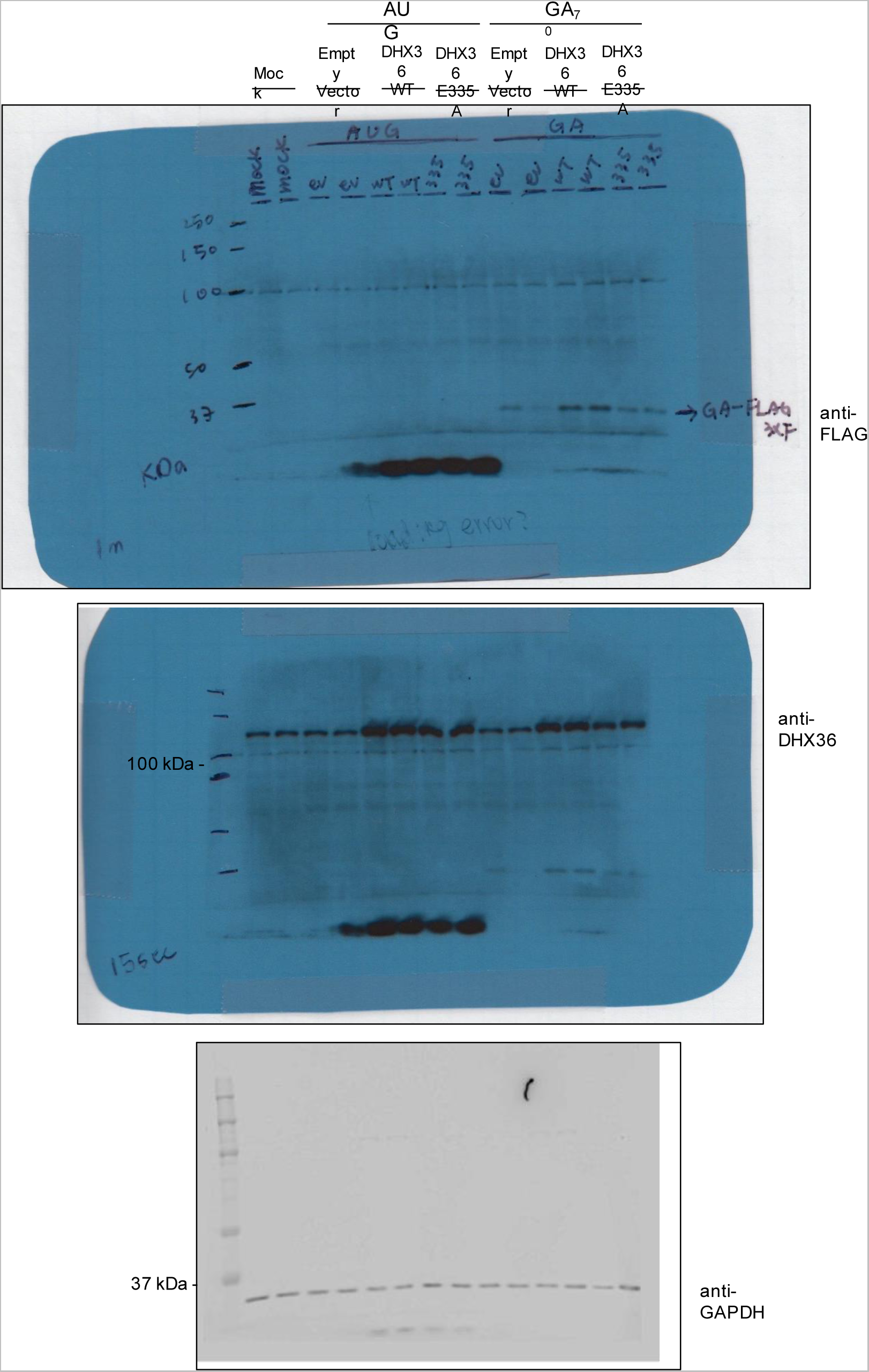

**Figure 6.**
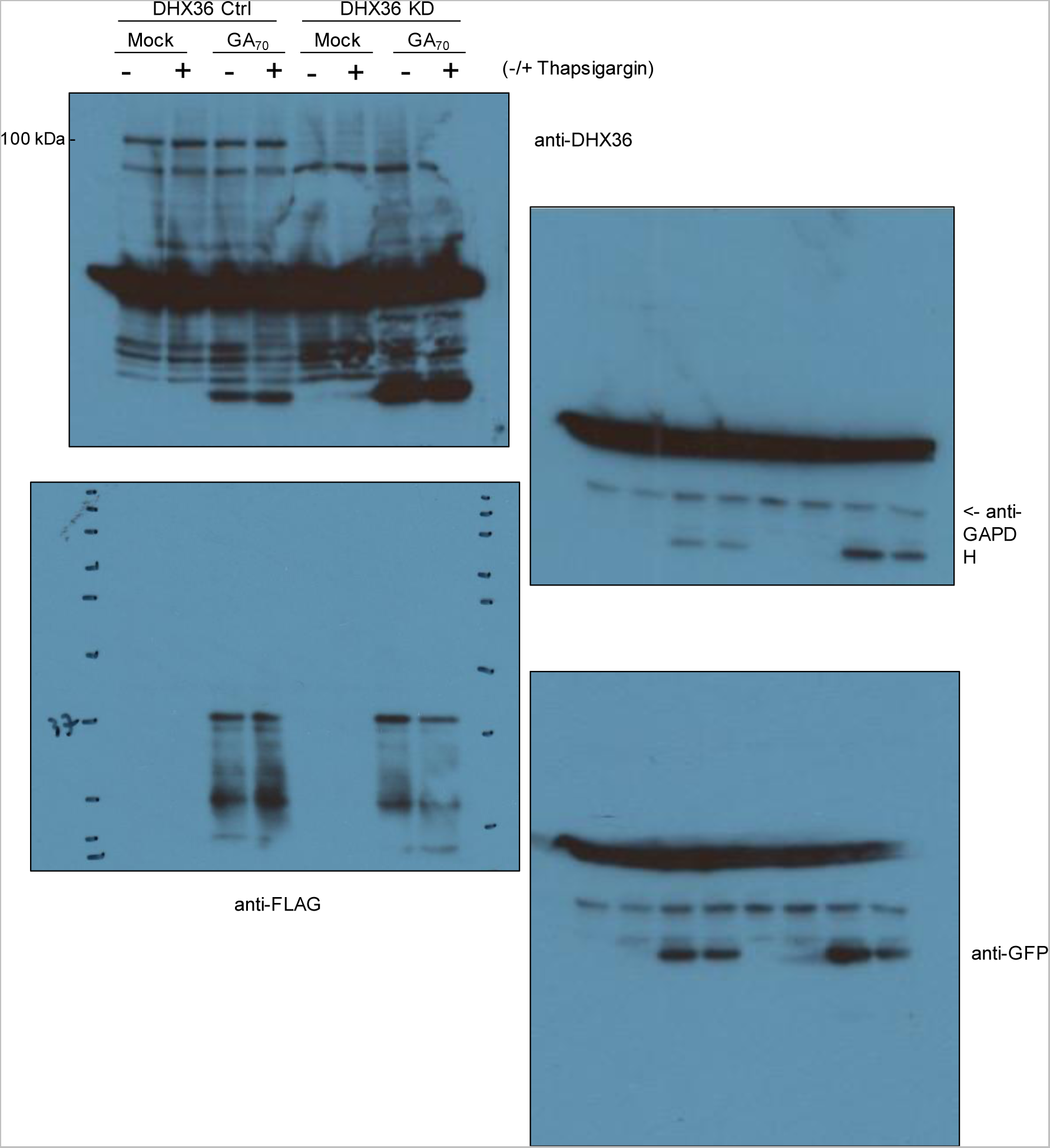

**Supplemental Figure 1.**
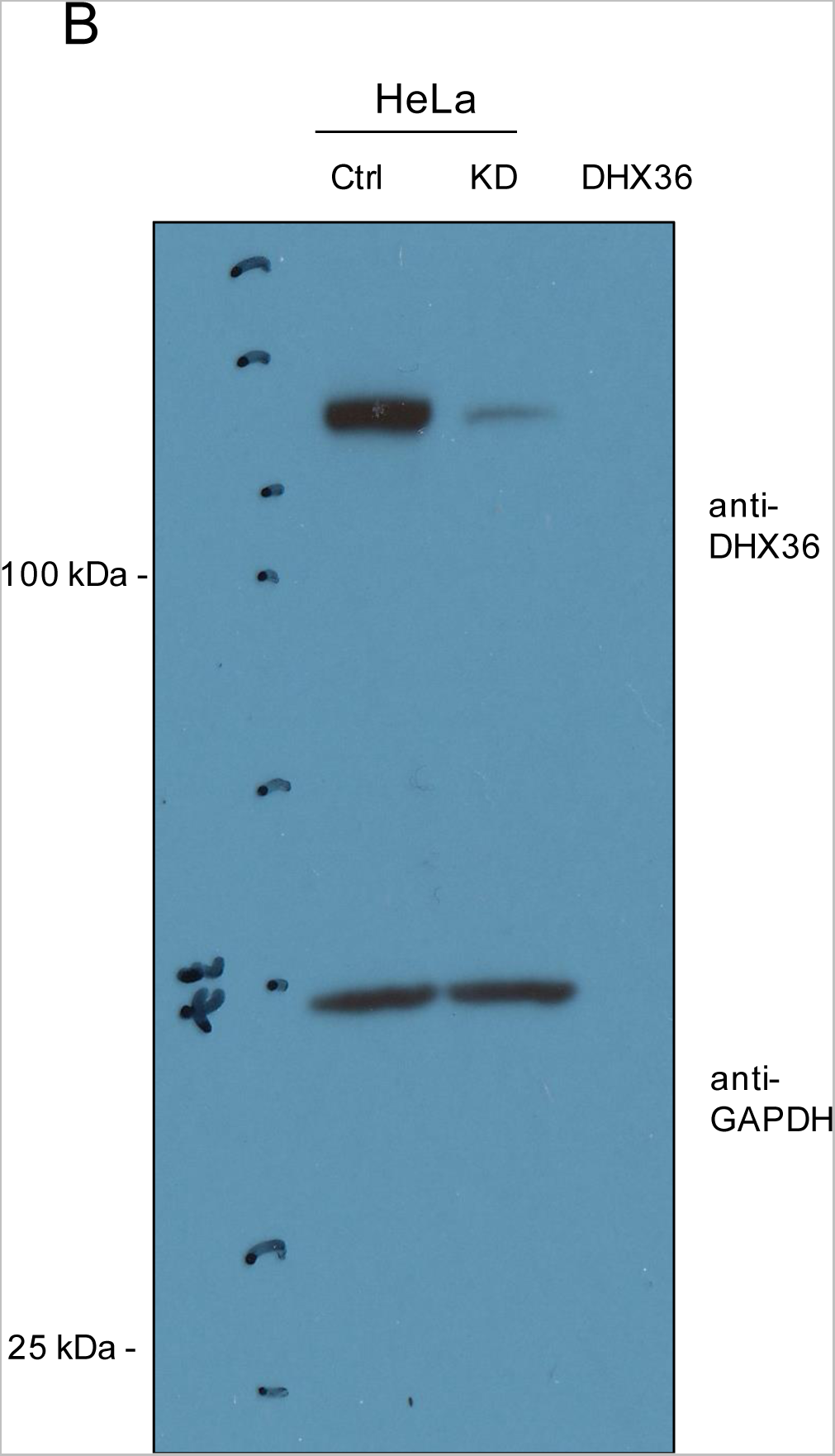
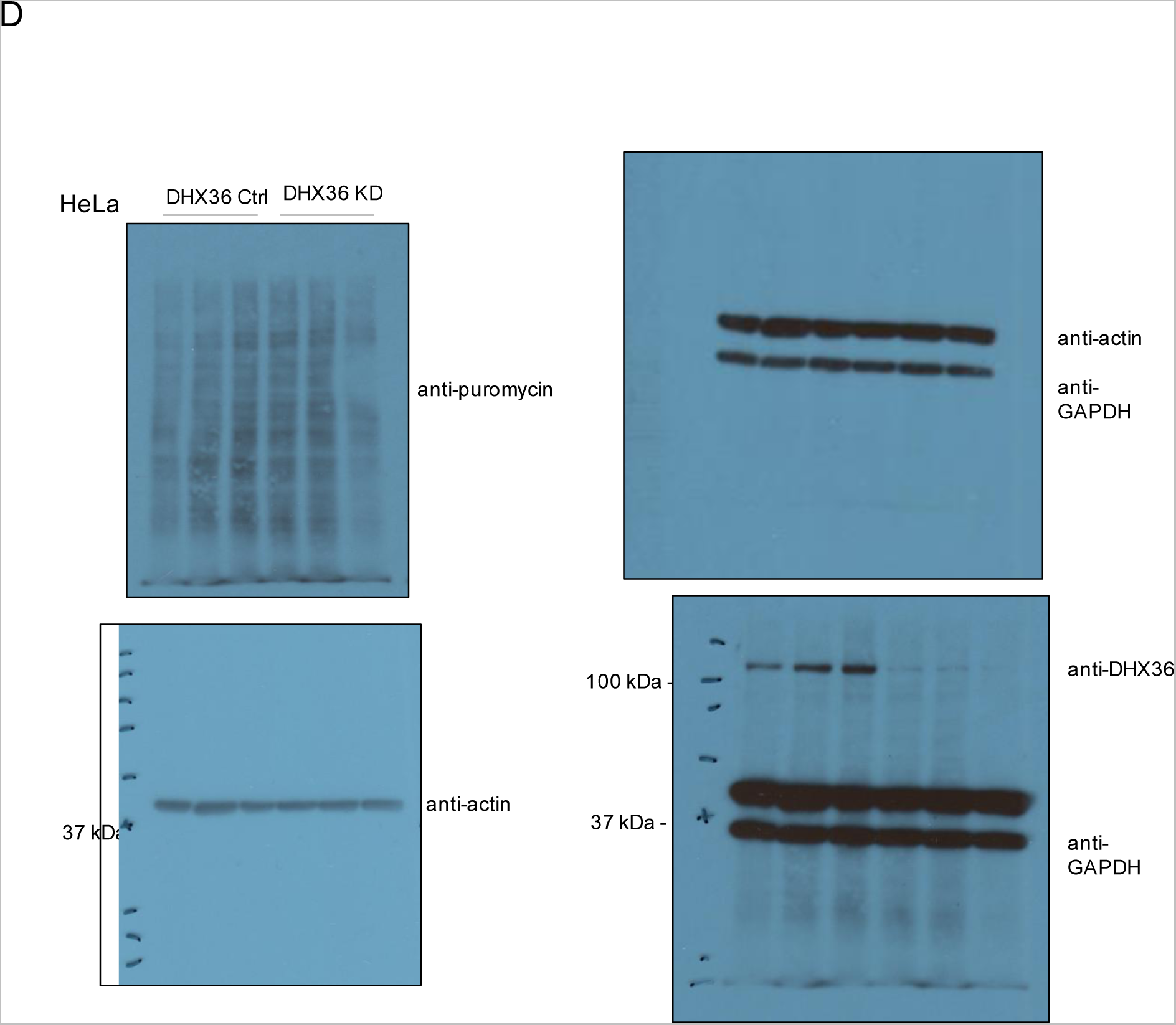

**Supplemental Figure 5.**
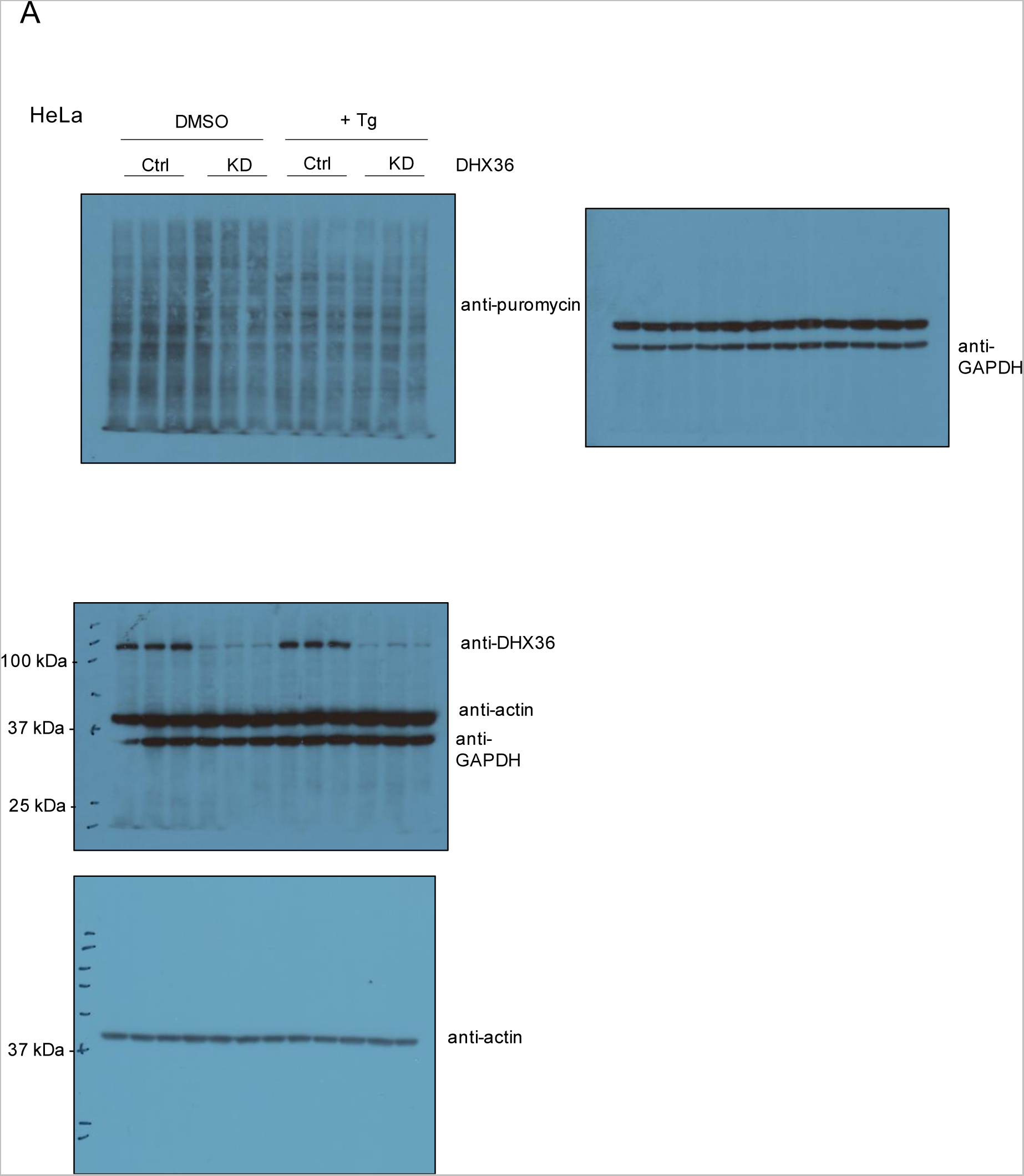

